# A direct RPA-Polθ interaction licenses theta-mediated end joining

**DOI:** 10.64898/2026.10.25.761560

**Authors:** Marco Barazas, Celeste I. Bos, Hanneke Kool, Joost Schimmel, Robin van Schendel, Martijn S. Luijsterburg, Marcel Tijsterman

**Author notes:** These authors contributed equally to this work (Marco Barazas, Celeste I. Bos).

## Abstract

DNA double-strand break repair often involves end resection, yet how resected ends are directed toward distinct repair pathways remains poorly understood. Here, we identify replication protein A (RPA) as an unexpected essential factor for cellular polymerase theta-mediated end joining (TMEJ). Using MUSIC-MAP, a newly developed platform linking endogenous sequence variants to DNA repair outcomes, we map a discrete functional interface between RPA2 and polymerase theta (Polθ). Disrupting this interface in RPA2 abolishes TMEJ without impairing cell viability or homologous recombination. Mutations in the complementary Polθ surface likewise impair TMEJ, whereas reciprocal charge-reversal mutations in RPA2 and Polθ restore TMEJ, providing genetic evidence for a direct physical interaction. Structural modeling suggests that the Polθ-binding surface becomes accessible at RPA-coated ssDNA termini, positioning RPA as a docking platform for Polθ engagement. These findings reveal how RPA, previously considered a barrier to TMEJ, instead functions as a molecular platform that translates the organization of ssDNA into selective enzyme engagement, enabling mutagenic repair without compromising homologous recombination.

## Introduction

DNA double-strand breaks (DSBs) are repaired by pathways with distinct substrate requirements, processing activities, and mutational consequences. Canonical non-homologous end joining (c-NHEJ) joins DNA ends with limited processing, whereas homologous recombination (HR) acts on resected ends and uses a homologous template to restore the damaged sequence(*1, 2*). Theta-mediated end joining (TMEJ) provides an alternative route for repairing resected DNA ends and depends on polymerase theta (Polθ, encoded by the gene *POLQ*), which contains an N-terminal helicase-like domain and a C-terminal DNA polymerase domain connected by a largely disordered central region (*3–5*). TMEJ uses short complementary sequences, termed microhomologies, to align DNA ends, followed by Polθ-mediated DNA synthesis that facilitates their joining. This generates microhomology-associated deletions, and, through iterative cycles of synthesis and release, templated insertions (*6–10*). These products provide a record of TMEJ activity(*11*).

TMEJ is particularly important in HR-deficient cells. Microhomology-associated deletions and templated insertions accumulate in BRCA1- and BRCA2-deficient tumors(*12, 13*), and loss or inhibition of Polθ is selectively toxic in HR-deficient backgrounds(*14–17*), motivating the development of Polθ inhibitors as candidate cancer therapeutics(*18, 19*). These findings implicate TMEJ in the repair of DNA intermediates that would otherwise be repaired by HR. TMEJ also operates in HR-proficient cells(*20*), including during mitosis, when HR and c-NHEJ are restricted(*21–26*). Its dependence on DNA-end resection requires Polθ to engage single-stranded DNA (ssDNA), which is rapidly coated upon formation by replication protein A (RPA), a heterotrimer composed of RPA70/RPA1, RPA32/RPA2, and RPA14/RPA3(*27*). Trimeric RPA contains four principal ssDNA-binding domains (DBDs): DBD-A, DBD-B and DBD-C in RPA1, and DBD-D in RPA2(*28*). RPA protects ssDNA and organizes protein recruitment during replication, recombination, checkpoint signaling, and repair(*29, 30*). RPA coating has generally been viewed as a barrier to TMEJ; the Polθ helicase-like domain can displace RPA to promote DNA annealing(*31–33*). How Polθ productively engages RPA-coated ssDNA to initiate TMEJ remains unclear. Here, we identify a direct interaction between RPA2 and Polθ that is essential for cellular TMEJ. By combining systematic genetic interrogation with structural analysis, we establish RPA as an active participant in TMEJ and propose a mechanism by which RPA facilitates Polθ engagement specifically at resected DNA ends.

## Results

### Genetic profiling identifies a TMEJ-specific surface in RPA2

We previously developed MUSIC (MUtational Scars of Induced DNA Cleavage), a genetic screening strategy that links gene perturbations to DSB-repair pathways by analyzing hundreds of thousands of repair junctions in bulk cell populations(*34, 35*). Starting from a genome-wide sgRNA screen in mouse embryonic stem (mES) cells, we identified an *Rpa2*-targeting sgRNA (sgRNA *Rpa2-4*) that produced a repair outcome profile closely resembling that caused by disruption of *Polq* (Fig. 1A): TMEJ-associated outcomes were strongly depleted, whereas c-NHEJ and homology-directed outcomes remained largely intact or increased. This repair phenotype was confirmed in a focused *Rpa2*-targeting sgRNA library (Fig. S1A-C), in which most *Rpa2*-targeting sgRNAs were depleted from the cell population, consistent with the essential functions of RPA2. In silico prediction of editing outcomes using inDelphi(*36*) identified two distinct in-frame 9 base-pair (bp) deletions among the predominant products of sg*Rpa2-4* (Fig. S1D), suggesting that specific alterations in this region might selectively compromise TMEJ.

**Figure 1.**
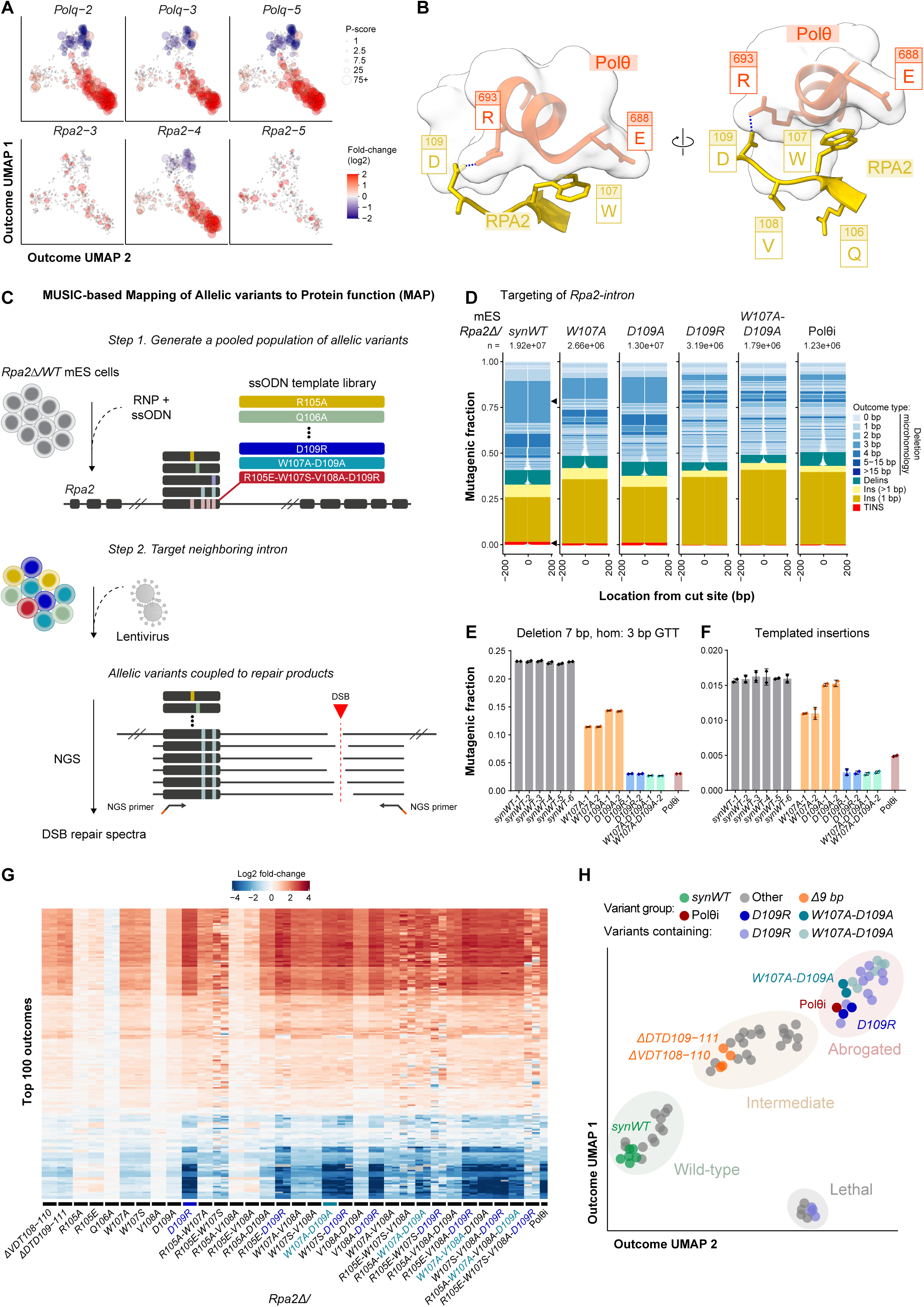
Genetic mapping of a TMEJ-specific surface in RPA2. (**A**) MUSIC-derived outcome UMAPs showing the effects of individual *Polq*- and *Rpa2*-targeting sgRNAs on DSB repair(*35*). Each point represents a distinct repair outcome, positioned according to similarities in its genetic dependencies. Colors indicate log2 fold-changes in outcome abundance relative to non-targeting controls, and point sizes reflect statistical significance (P-score). (**B**) AlphaFold3-predicted interaction between RPA2 (mouse, yellow) and the helicase-like domain of Polθ (mouse, red). The predicted interface involves RPA2 residues W107 and D109 and a complementary surface in Polθ, including E688 and R693. (**C**) Schematic overview of MUSIC-MAP (MUSIC-based Mapping of Allelic variants to Protein function). First, hemizygous *Rpa2* mES cells are transfected with Cas9 ribonucleoprotein (RNP) complexes and a library of single-stranded oligodeoxynucleotide (ssODN) repair templates to generate a pooled population of cells carrying defined allelic variants. Subsequently, a nearby intronic site is targeted using lentiviral delivery of a Cas9 sgRNA to induce DSBs. The allelic variant and associated repair outcome are captured within the same sequencing fragment, allowing direct assignment of repair products to the genotype in which they were generated. For simplicity, only one allelic variant is illustrated, showing how multiple DSB repair outcomes are generated and collectively define its repair profile. In practice, each allelic variant is represented by numerous independently engineered cells. **(D)** Representative DSB repair outcome spectra obtained by MUSIC-MAP and analyzed using SIQ software (*1, 2*). Tornado plots display individual repair products according to their frequency among mutagenic reads (y-axis) and position relative to the cleavage site (x-axis). Outcomes are colored by repair product type, with white regions indicating deleted sequences. Profiles are shown for synonymously encoded wild-type controls (synWT), the indicated RPA2 substitutions, and cells treated with the Polθ inhibitor ART558 (10 µM; Polθi). Data from two biological replicates were combined. A recurrent 11 bp deletion involving 3 bp microhomology and templated insertions, depleted upon Polθ inhibition, are highlighted (black arrowhead). Delins, deletion with insertion; Ins, insertion; TINS, templated insertion; bp, base pair. (**E,F**) Quantifications of the recurrent 11 bp deletion with 3 bp microhomology (E) and templated insertions (F) across the indicated RPA2 variants and Polθ-inhibited cells. The outcome frequency among mutagenic reads is plotted for individual synonymous encodings of each variant. Bars represent mean ± SD, and diamonds indicate measurements from two independent biological replicates (independent bulk RNP transfections). Hom, homology. (**G**) Heatmap of MUSIC-MAP-derived repair profiles for the indicated *Rpa2* allelic variants. Log2 fold-changes in the frequencies of the 100 most abundant repair outcomes relative to synonymously encoded wild-type controls are shown. Individual synonymous encodings and biological replicates are displayed separately, with variants grouped by amino acid substitution. Repair outcomes are ordered by unsupervised hierarchical clustering. Grey indicates outcomes for which no supporting reads were detected. (**H**) UMAP representation of MUSIC-MAP repair profiles, positioning individual allelic variants according to similarities in their repair outcome distributions. Synonymous encodings are displayed separately. Variants carrying D109R or W107A-D109A substitutions are highlighted, together with synonymous wild-type controls, Polθ-inhibited cells, and variants carrying 9 bp deletions. The analysis reveals distinct groups of variants with wild-type-like, intermediate, or strongly impaired TMEJ activity, as well as a separate cluster potentially associated with impaired cell viability.

Independent introduction of one of these deletions using an alternative sgRNA reproduced the repair phenotype, supporting an on-target effect (Fig. S1E). AlphaFold3(*37*) modeling placed this region, encompassing RPA2 residues 106–109, within the ssDNA-binding domain D (DBD-D), against a surface in the Polθ N-terminal helicase domain (Fig. 1B and Fig. S2). The predicted interface involved both hydrophobic and electrostatic contacts, including RPA2 W107, which packs against a Polθ surface formed by E688, G689 and A692. Nearby, RPA2 D109 forms a putative salt bridge with Polθ R693. Together, these observations suggested a candidate RPA2–Polθ interface involved in TMEJ.

To systematically test this interface, we developed and applied MUSIC-MAP (MUSIC-based Mapping of Allelic variants to Protein function), which couples variants engineered at an endogenous locus to their DSB-repair spectra in pooled cell populations (Fig. 1C). In mES cells, modified to carry a single functional *Rpa2* allele (Fig. S3A), a library of 70 single-stranded donor oligonucleotides introduced alanine and polarity- and charge-reversal substitutions, combinations of substitutions at predicted contact residues, and substitutions at residues oriented away from the interface predicted by AlphaFold3. Each variant was endogenously encoded by two synonymous donor sequences, and six synonymous wild-type alleles served as internal controls. The edited population was then targeted with an sgRNA directed against an intron neighboring the exon encoding the interface. Sequencing across both sites captured the engineered *Rpa2* allele and the repair outcome in the same read, enabling repair profiles to be assigned to individual variants (Fig. 1C). Polθ inhibition with ART558 provided a TMEJ-deficient reference profile(*38*). MUSIC-MAP thereby enabled systematic functional profiling of endogenous alleles expressed at physiological levels by aggregating repair outcomes across independently edited cells.

The robustness of MUSIC-MAP was supported by wild-type repair profiles from synonymous variants, concordant results from independent donor encodings of identical amino acid substitutions and between replicates, and recapitulation of the initial sgRNA-screen phenotype by engineered 9 bp deletions (Fig. S4A-C). Substitutions at residues predicted to face away from Polθ, including Q106 and V108 produced wild-type repair profiles (Fig. S4A). In contrast, individual alanine substitutions at W107 or D109 partially reduced TMEJ, whereas the W107A-D109A double substitution produced a repair profile closely resembling that induced by Polθ inhibition (Fig. 1D-F). Most strikingly, a single charge-reversal substitution, D109R, abolished TMEJ-associated repair products.

Across the repair spectrum, interface-disrupting RPA2 variants recapitulated the pattern of outcome changes induced by Polθ inhibition (Fig. 1G). Consistent with this, UMAP analysis placed D109R and W107A-D109A near the Polθ-inhibited reference, whereas variants with partial TMEJ defects occupied intermediate positions (Fig. 1H, S4D). Some combinatorial variants, including those containing R105E-W107S yielded markedly fewer repair outcomes (Fig. S4A), potentially reflecting broader impairment of RPA functions. Both R105 and W107 are predicted to contact ssDNA when DBD-D engages ssDNA, providing a possible explanation for the phenotype of the double substitution. By contrast, D109 is not predicted to contact ssDNA in this configuration, yet its charge reversal abolished TMEJ. We therefore focus subsequent analyses on D109 to investigate the functional importance of the predicted electrostatic interaction with Polθ.

Together, these findings identify a discrete surface in RPA2 that is critical for TMEJ and suggest that its contribution to this repair pathway can be separated from RPA’s canonical role in ssDNA binding.

### The RPA2 interface selectively controls TMEJ

We next tested whether the phenotype of the MUSIC-MAP-identified *Rpa2* allele variants persisted in endogenous *Rpa2 W107A-D109A* or *D109R* alleles, disrupting the predicted RPA2-Polθ interface through alanine substitutions or charge reversal, respectively (Fig. 2A, S3C). Both genotypes were recovered readily, and proliferation assays detected no differences in cell cycle distribution or growth rate relative to wild-type cells (Fig. 2B-C), indicating that the interface mutations preserve the essential cellular functions of RPA2. Nevertheless, repair profiling at two genomic sites confirmed an almost complete loss of TMEJ-associated outcomes in both mutant cells (Fig. 2D-I): microhomology-associated deletions (Fig. 2E, 2H) and templated insertions (Fig. 2F, 2I) were depleted to an extent comparable to that observed after *Polq* knockout. As an independent functional readout of TMEJ, we examined sensitivity to ionizing radiation (IR)(*14, 20*). Both *Rpa2*-interface mutants were hypersensitive to IR, with sensitivities matching that of *Polq*-deficient mES cells (Fig. 2J). Polθ inhibition sensitized wild-type cells but did not further sensitize either *Rpa2* mutant (Fig. 2K), consistent with shared pathway defects.

**Figure 2.**
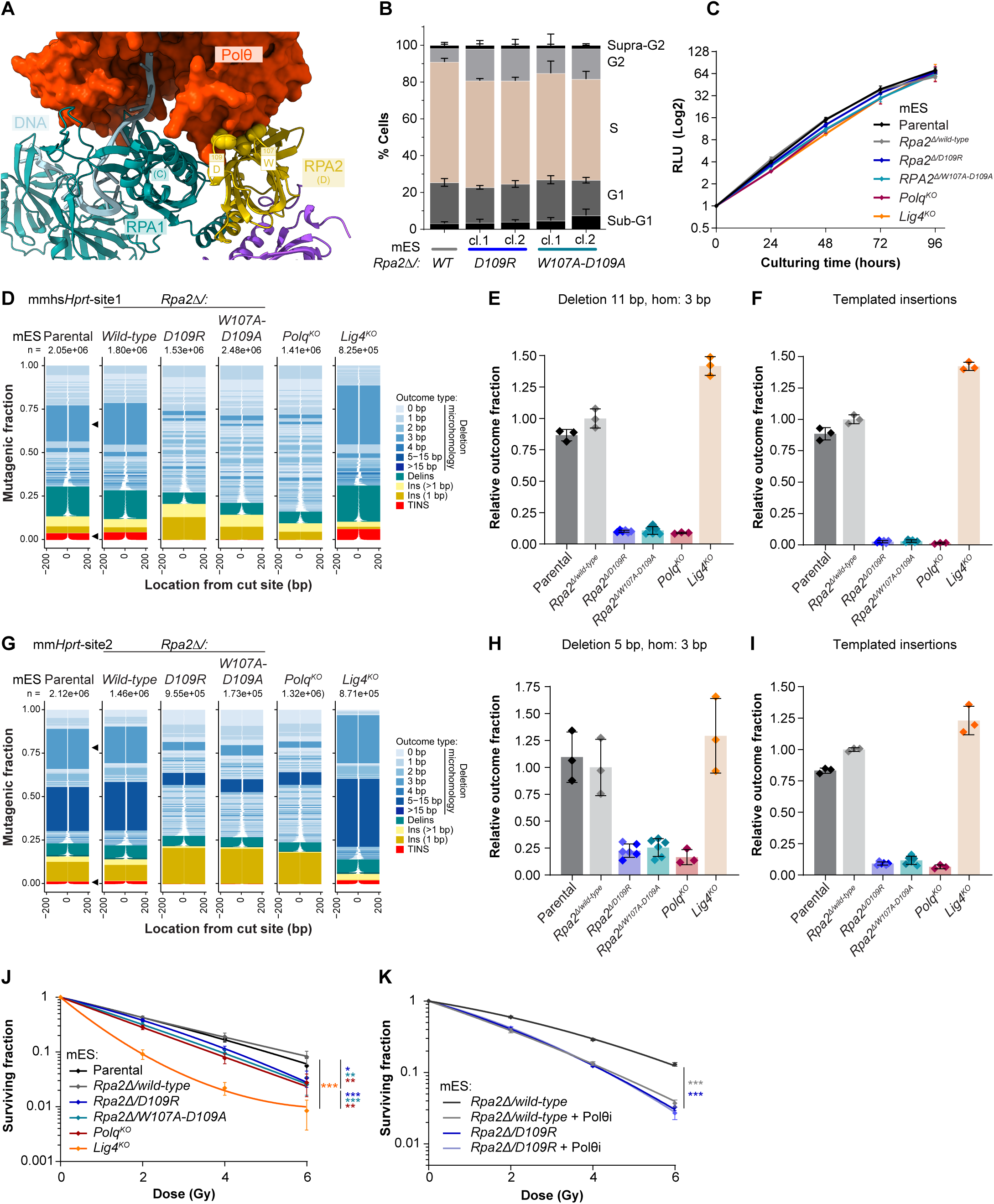
The RPA2 interface selectively controls TMEJ. (**A**) AlphaFold3-predicted model of the RPA heterotrimer (mouse) bound to ssDNA and interacting with Polθ (mouse). The DNA-binding domains RPA1 DBD-C and RPA2 DBD-D are indicated, together with RPA2 residues W107 and D109 that form part of the predicted Polθ-binding interface. (**B**) Cell cycle distributions of wild-type and *Rpa2* knock-in mutant mES cells determined by propidium iodide (PI) staining. Cells predominantly reside in S phase, with comparable cell cycle distributions across genotypes. Data are presented as mean ± SD from three independent experiments. WT, wild-type; cl., clone. (**C**) Growth curves of the indicated mES cell lines showing comparable proliferation rates. Measurements from technical duplicates were averaged, normalized to the initial time point, and log2-transformed. Data are presented as mean ± SEM from three independent biological replicates. **(D, G)** DSB repair outcome spectra at two endogenous *Hprt* target sites (site 1 and site 2, respectively) in mES cells of the indicated genotypes. Repair outcomes were identified from short-amplicon Illumina sequencing data using SIQ and visualized with SIQplotter (2). Tornado plots show the distribution of individual repair outcomes among mutagenic reads, with outcomes positioned relative to the cleavage site and colored according to repair product type. White regions indicate deleted sequences. Data from three biological replicates were averaged (fractions), and repair outcomes are displayed proportionally according to their relative abundance. For figure clarity, data from two independently derived *Rpa2 D109R* and *Rpa2 W107A-D109A* clones were combined. Recurrent microhomology-mediated deletions and templated insertions are indicated (black arrowheads). Delins, deletion with insertion; Ins, insertion; TINS, templated insertion; bp, base pair. **(E, H)** Quantification of recurrent microhomology-mediated deletions at *Hprt* site 1 (11 bp deletion with 3 bp microhomology) and site 2 (5 bp deletion with 3 bp microhomology), respectively, indicated by black arrowheads in (D) and (G). Data from two independently derived *Rpa2 D109R* and *Rpa2 W107A-D109A* clones (differently shaded) were combined. Hom, homology. **(F, I)** Quantification of templated insertions at *Hprt* sites 1 and 2, respectively. For (E, F, H, I), relative outcome fractions were normalized to the parental wild-type control. Individual data points represent biological replicates (n = 3), and bars indicate mean ± SD. Hom, microhomology. Data from two independently derived *Rpa2 D109R* and *Rpa2 W107A-D109A* clones (differently shaded) were combined. Hom, homology. (**J**) Ionizing radiation (IR) sensitivity of the indicated mES cell lines determined by clonogenic survival. Data from two independently derived RPA2 D109R and RPA2 W107A-D109A clones were combined for analysis and presentation. Data are shown as mean ± SEM from three independent biological replicates. Survival curves were fitted using a linear-quadratic model, and differences between model fits were assessed by ANOVA followed by Benjamini-Hochberg correction for multiple testing. Significant comparisons were *Rpa2Δ/WT* versus *Lig4^KO^* (*P* = 1.46 × 10⁻⁵), *Rpa2Δ/WT* versus *Rpa2Δ/D109R* (*P* = 0.0324), *Rpa2Δ/WT* versus *Rpa2Δ/W107A-D109A* (*P* = 0.00909), *Rpa2Δ/WT* versus *Polq^KO^* (*P* = 0.00909), *Rpa2Δ/D109R* versus *Lig4^KO^* (*P* = 1.22 × 10⁻⁵), *Rpa2Δ/W107A-D109A* versus *Lig4^KO^* (*P* = 0.000159), and *Polq^KO^* versus *Lig4^KO^* (*P* = 0.00295). (**K**) Epistasis analysis of RPA2 D109R and Polθ inhibition in response to IR, assessed by clonogenic survival in the presence or absence of the Polθ inhibitor ART558 (10 µM). Data from two independently derived RPA2 D109R clones were combined for analysis and presentation. Data are shown as mean ± SEM from three independent biological replicates. Survival curves were fitted using a linear-quadratic model, and differences between model fits were assessed by ANOVA followed by Benjamini-Hochberg correction for multiple testing. Significant comparisons were *Rpa2Δ/WT* versus *Rpa2Δ/WT* + Polθi (*P* = 8 × 10⁻⁸) and *Rpa2Δ/WT* versus *Rpa2Δ/D109R* (*P* = 1.6 × 10⁻¹³).

### The RPA2-Polθ interface is conserved and dispensable for homologous recombination

To assess whether disruption of the RPA2-Polθ interface selectively affects TMEJ or also compromises other RPA-dependent repair functions, we examined HR-associated responses. *Rpa2 D109R* and *Rpa2 W107A-D109A* cells showed no detectable changes in IR-induced RAD51 focus formation or sensitivity to PARP inhibition (Fig. 3A-B, S5), supporting preservation of HR-associated DNA damage responses(*39*).

**Figure 3.**
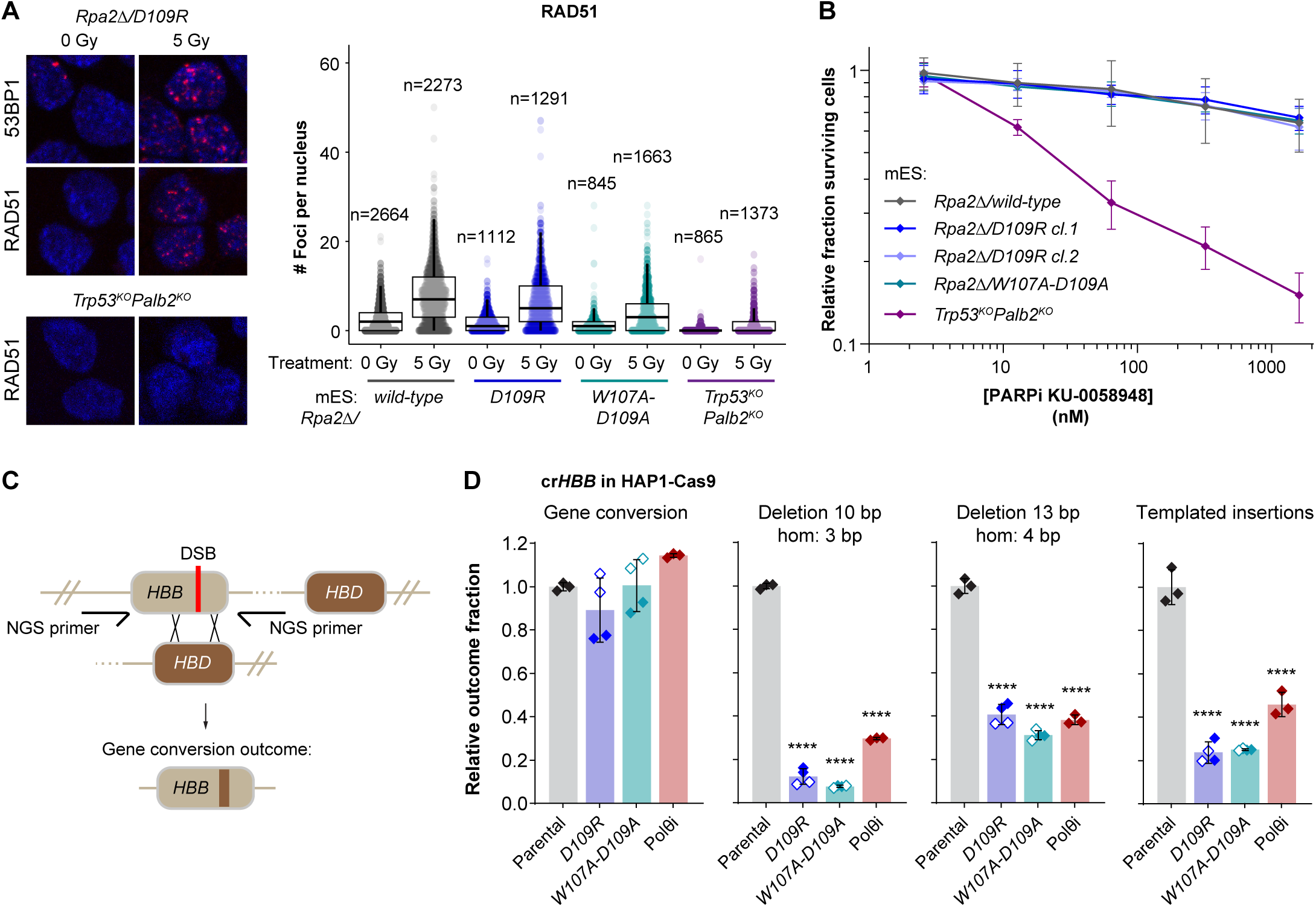
The RPA2-Polθ interface is dispensable for homologous recombination. (**A**) Analysis of ionizing radiation (IR)-induced RAD51 foci formation in mES cells of the indicated genotypes. Cells were fixed 3 hours after irradiation (5 Gy), stained for 53BP1 or RAD51, and imaged by confocal microscopy. Representative images and quantification of RAD51 foci per nucleus are shown. At least four fields were imaged per genotype and biological replicate, and the experiment was performed at least three times independently. Box-and-whisker plots show the distribution of foci counts across individual nuclei pooled from all experiments, with the number of nuclei analyzed indicated above each group. Statistical analyses were performed on biological replicate means (see also Fig. S5). (**B**) PARP inhibitor (PARPi) sensitivity of mES cells of the indicated genotypes. Relative cell survival was determined from cell counts after 96 hours of treatment with the PARP inhibitor KU-0058948. Measurements from technical triplicates were averaged and normalized to untreated controls. Data are presented as mean ± SEM from three independent biological replicates. (**C**) Schematic representation of the *HBB*-*HBD* gene conversion assay. Following Cas9-induced cleavage at the *HBB* locus, homologous sequences in *HBD* can serve as a repair template. Gene conversion events using *HBD* as a donor can be identified by the incorporation of *HBD*-specific sequence variants into the repaired *HBB* locus, allowing these outcomes to be distinguished from other DSB repair products by sequencing. (**D**) Quantification of DSB repair outcomes following Cas9-induced cleavage at the *HBB* locus in HAP1-Cas9 cells of the indicated genotypes. Four outcome classes were analyzed: *HBD*-templated gene conversion, a 10 bp deletion with 3 bp microhomology, a 13 bp deletion with 4 bp microhomology, and templated insertions. Relative outcome fractions were normalized to the parental control. Data represent independent biological replicates (n = 3 for parental controls and n = 2 per independently derived *RPA2* mutant clone), with two independently derived clones analyzed for each RPA2 mutant genotype. Data are presented as mean ± SD. Statistical significance was assessed by ordinary one-way ANOVA followed by Dunnett’s multiple-comparisons test, comparing each condition with the parental control. Differences in gene conversion frequencies were not significant. \*\*\*\**P* < 0.0001 (adjusted).

We next extended our analysis to human cells to test evolutionary conservation and directly measure homology-directed gene conversion using an established assay(*40*). We generated monoclonal HAP1-Cas9 lines carrying endogenous *RPA2 D109R* or *W107A-D109A* substitutions (Fig S3D). At the *HBB* locus, a Cas9-induced DSB can undergo gene conversion using the homologous *HBD* sequence, generating sequence changes distinguishable from end-joining outcomes (Fig. 3C). Both mutants retained gene-conversion frequencies comparable to those of parental cells (Fig. 3D). In the same assay, microhomology-associated deletions and templated insertions were markedly reduced (Fig. 3E-F), reproducing the observed loss of TMEJ-associated outcomes in mES cells. Repair profiling at a second target, *HPRT*, likewise showed a pronounced reduction in templated insertions and a selected (Polθ-dependent) microhomology-associated deletion (Fig. S6); Polθ inhibition reduced the same repair products in parental HAP1 cells.

Collectively, these experiments establish that the RPA2 interface is conserved between mouse and human and is specifically required for cellular TMEJ while leaving the essential and recombinational functions of RPA intact.

### A complementary surface in Polθ is required for TMEJ

Having identified the RPA2 residues required for TMEJ, we next examined the complementary surface in Polθ predicted by AlphaFold3(*37*). The model positioned a surface within the N-terminal helicase-like domain of Polθ against RPA2 residues 105-109 (Fig. 4A). Polθ residues E688, G689 and A692 (mouse numbering) form a surface that packs against RPA2 W107. Nearby, Polθ R693 forms a putative salt bridge with RPA2 D109, while K696 and K698 may provide additional electrostatic contacts across the interface.

**Figure 4.**
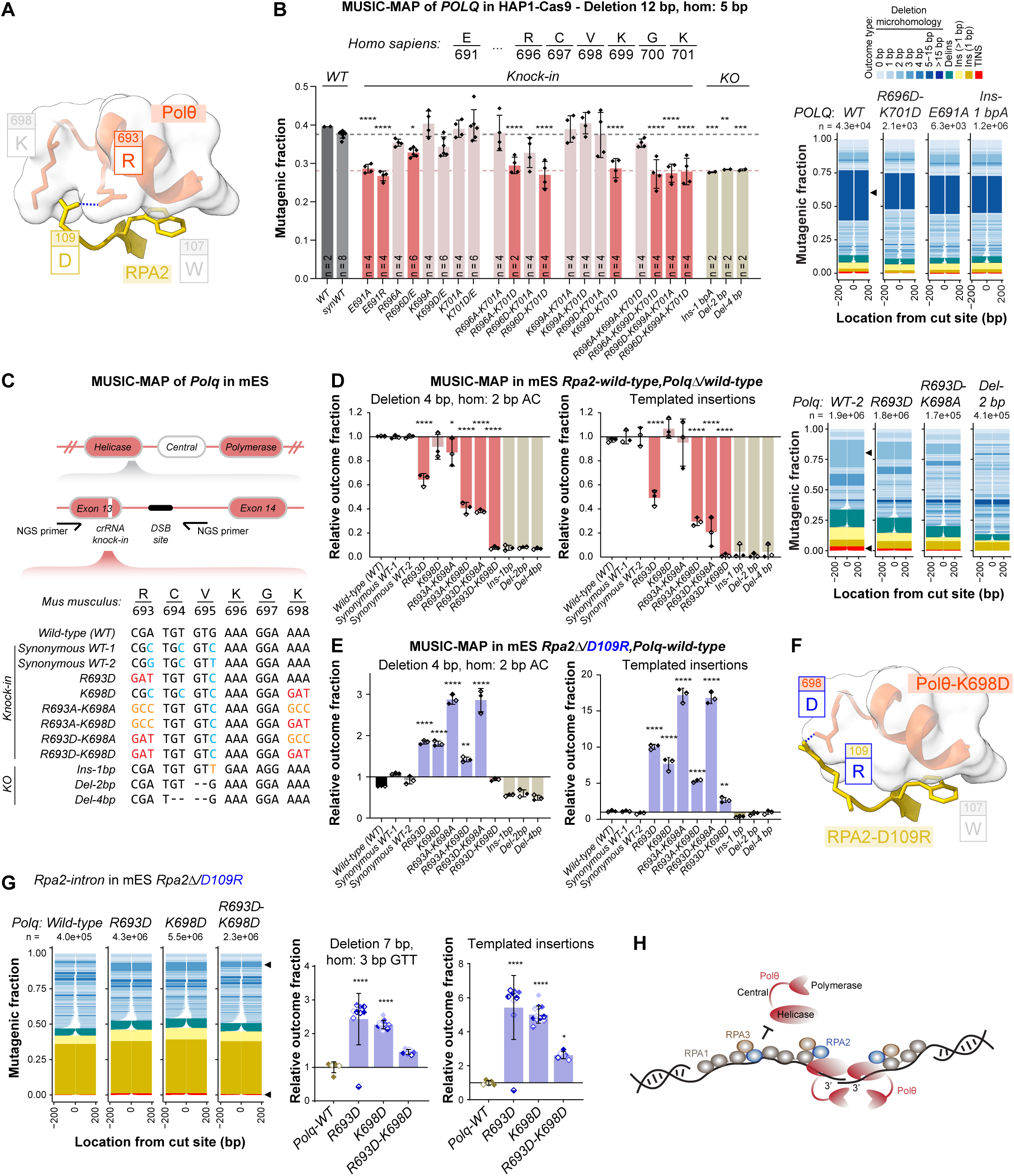
Mapping of the complementary surface in Polθ and genetic rescue of RPA2-D109R. (**A**) AlphaFold3-predicted interaction between RPA2 (mouse) and the helicase-like domain of Polθ (mouse), highlighting the predicted electrostatic interaction between RPA2 D109 and Polθ R693, with Polθ K698 positioned nearby. (**B**) MUSIC-MAP analysis of the complementary RPA2-interacting surface in human *POLQ*, performed in HAP1-Cas9 cells using single and combinatorial substitutions at residues E691, R696, K699, and K701. Sequencing reads were assigned to individual allelic variants, and associated DSB repair outcome profiles were generated using SIQ software. Tornado plots display individual repair products according to their frequency among mutagenic reads (y-axis) and position relative to the cleavage site (x-axis), data from two biological replicates and synonymous variant encodings were combined (fractions). A recurrent 12 bp deletion involving 5 bp microhomology, depleted in *POLQ* frameshift alleles, was selected for quantification (black arrowhead). Bars represent mean ± SD. Measurements from two independent pooled RNP transfections and two synonymous encodings per variant were combined for presentation. Only the three most abundant frameshift alleles are shown. Statistical significance was assessed by one-way ANOVA followed by Dunnett’s multiple-comparisons test, using synonymously encoded wild-type controls as the reference. \**P* < 0.05, \*\**P* < 0.005, \*\*\**P* < 0.0005, \*\*\*\**P* < 0.0001. Delins, deletion with insertion; Ins, insertion; TINS, templated insertion; bp, base pair; hom, homology. (**C**) Schematic overview of the focused MUSIC-MAP strategy targeting the endogenous *Polq* locus in mES cells. Defined allelic variants were introduced into the region encoding the predicted RPA2-interacting surface using ssODN templates, followed by DSB induction at a nearby genomic site to generate allele-associated repair profiles. The wild-type sequence, synonymous wild-type encodings, engineered substitutions, and representative frameshift alleles are shown. (**D**) MUSIC-MAP analysis of *Polq* allelic variants in mES cells hemizygous for *Polq* and expressing wild-type RPA2. Sequencing reads were assigned to individual allelic variants, and repair outcomes were identified using SIQ and visualized with SIQplotter. Representative repair spectra are shown for the indicated variants (three biological replicates combined). Color legend as in (B). A recurrent 4 bp deletion involving 2 bp microhomology and templated insertions (black arrowheads), both dependent on Polθ activity as established by their depletion in *Polq* frameshift alleles, were quantified. Outcome fractions were normalized to the mean of synonymously encoded wild-type controls and are presented as mean ± SD from three independent biological replicates (pooled RNP transfections). Statistical analyses were performed as in (B), using synonymous WT-1 as the reference. (**E**) MUSIC-MAP analysis of *Polq* allelic variants, performed as in (D), in mES cells carrying endogenous RPA2 D109R. Complementary charge-reversal substitutions in Polθ restore TMEJ-associated repair outcomes impaired by RPA2 D109R, including the recurrent microhomology-mediated deletion and templated insertions. Data are presented and analyzed as in (D), allowing direct comparison of variant-specific effects between RPA2 wild-type and D109R backgrounds. (**F**) AlphaFold3 model of the RPA2 D109R-Polθ K698D interface, illustrating a predicted salt bridge formed between the charge-reversed residues. The D109R substitution is predicted to reposition the arginine side chain toward Polθ residue 698, potentially enabling an electrostatic interaction with K698D that is absent between the corresponding wild-type residues. (**G**) Validation of genetic rescue in independently derived monoclonal mES cell lines carrying the indicated *Polq* substitutions in the RPA2 D109R background. DSB repair was profiled at the *Rpa2* locus near the engineered D109R substitution. Repair profiles from independent clones of the same genotype were combined for visualization, and the total number of mutagenic reads is indicated above each plot. A recurrent 7 bp deletion involving 3 bp microhomology and templated insertions (black arrowheads) were quantified. Outcome fractions were normalized to the *Polq* wild-type control and are presented as mean ± SD from independently generated clones. Statistical analyses were performed as in (B), using *Polq* wild-type as the reference. (**H**) Schematic model illustrating how the interaction between RPA2 and Polθ may facilitate engagement of Polθ at resected DNA ends. Within RPA-coated ssDNA (based on (*29*)), binding of RPA2 DBD-D to DNA may restrict accessibility of the Polθ-interacting surface. At ssDNA termini, limited DNA availability or transient disengagement of DBD-D could expose this interface, allowing Polθ to engage RPA2 residues W107 and D109 while RPA remains associated with DNA. This interaction may position the Polθ helicase-like domain to engage ssDNA and facilitate its transfer toward the polymerase domain. RPA1 DNA-binding domains A–C, RPA2 DBD-D, and RPA3 DBD-E are represented schematically, together with the Polθ helicase-like and polymerase domains connected by a central region

To genetically define this interface, we applied MUSIC-MAP in human HAP1 cells to test substitutions at these and neighboring residues, individually and in combination. To assess TMEJ activity, we profiled DSB repair at an endogenous target site near the introduced *POLQ* mutations, focusing on a recurrent 12 base-pair deletion with 5 base-pair microhomology that was reduced in cells carrying *POLQ* frameshift alleles (Fig. 4B). Effects on TMEJ depended on both the residue and the substitution. Whereas several single substitutions retained near-wild-type activity, Polθ E691 (E688 in mouse) and a charge reversal at R696 (R696D; R693D in mouse) reduced TMEJ, providing further support for the AlphaFold3 model: E691 in Polθ packs against the aromatic ring of W107 in RPA2, while K699 further stabilizes RPA2 D109 against Polθ R696. Combining substitutions at R696 and K701 further impaired TMEJ activity, with R696D-K701D producing reductions comparable to those observed for *POLQ* knockout alleles. Together, these results establish the functional importance of the predicted RPA2-binding surface of Polθ for TMEJ.

We next examined the corresponding residues in mES cells with a focused MUSIC-MAP library containing synonymous wild-type controls, single charge reversals, and combinations of alanine and charge-reversal substitutions (Fig. 4C). Synonymous controls retained wild-type repair profiles. Charge reversal at R693 (R693D) reduced both a recurrent microhomology-associated deletion and templated insertions, whereas K698D alone had little effect on either readout (Fig. 4D). Combining R693D with substitution of K698 further reduced both classes of repair products, in line with the prediction that K698 provides additional electrostatic contacts across the RPA2-Polθ interface.

These analyses identified a conserved functional surface in the Polθ helicase-like domain, with the position- and substitution-dependent effects suggesting contributions from multiple contacts, including the predicted interaction between RPA2 D109 and Polθ R693 (R696 in human).

### Complementary charge reversal restores TMEJ

We next asked whether TMEJ could be restored by re-establishing electrostatic complementarity between RPA2 and Polθ. In the predicted interface, RPA2 D109 is positioned near the positively charged Polθ residues R693 and K696, while the neighboring K698 provides additional contacts across the interface. Reversing the charge of D109 to arginine would therefore be expected to disrupt these interactions, potentially introducing electrostatic repulsion. We reasoned that complementary charge reversals in Polθ, replacing R693 or K698 with negatively charged aspartate, might restore favorable interactions with RPA2 D109R, despite disrupting the interface with wild-type RPA2. To test this prediction, we applied the focused *Polq* MUSIC-MAP library in *Rpa2 D109R* mES cells, allowing the effects of identical *Polq* substitutions to be compared in wild-type and mutant *Rpa2* backgrounds (Fig. 4D,E).

The effects of *Polq* substitutions were strikingly dependent on the *Rpa2* genotype. Whereas Polθ R693D impaired TMEJ in cells expressing wild-type RPA2 (Fig. 4D), it restored both microhomology-associated deletions and templated insertions in *Rpa2 D109R* cells (Fig. 4E). Similarly, Polθ K698D rescued these repair outcomes in the *Rpa2 D109R* background, despite having little effect in wild-type cells. AlphaFold3 modeling provided a structural rationale for the latter rescue: the D109R substitution is predicted to reposition the arginine side chain toward Polθ residue 698, potentially enabling a new salt bridge with K698D that is absent between the corresponding wild-type residues (Fig. 4F). Several additional combinations containing a charge reversal at either position also increased TMEJ-associated outcomes, whereas synonymous wild-type did not impact TMEJ. Notably, combining R693D and K698D was less effective than either substitution alone, indicating that rescue depends on the precise electrostatic configuration of the interface rather than simply on the introduction of negative charge. Together, these results provide genetic evidence that complementary charge reversals across the predicted RPA2-Polθ interface can restore TMEJ.

To validate the rescue observed by MUSIC-MAP, we generated independent *Rpa2 D109R* mES clones carrying *Polq R693D*, *Polq K698D*, or the combined *R693D-K698D* substitutions (Fig. S3E) and profiled DSB repair at three genomic targets: two sites in *Hprt* and one in *Rpa2* (Fig. 4G, S7). At all three sites, either single Polθ charge reversal increased both microhomology-associated deletions and templated insertions relative to *Rpa2 D109R* cells retaining wild-type Polθ. The combined *R693D-K698D* substitution produced a smaller increase in both repair outcomes, confirming the configuration-dependent rescue observed by MUSIC-MAP. Thus, complementary *Polq* mutations reproducibly suppressed the RPA2 D109R repair defect in independently derived clones and across distinct genomic substrates.

The reversal of the functional effect of Polθ R693D, from impairing TMEJ with wild-type RPA2 to restoring it with RPA2 D109R, provides strong genetic evidence that TMEJ depends on the compatibility of the two protein surfaces. Together with the reciprocal interface maps, these findings establish the functional importance of the predicted direct RPA2-Polθ interaction.

### Structural modeling suggests how Polθ engages RPA-coated DNA

The functional maps closely matched the RPA2-Polθ interface predicted by AlphaFold3(*37*) (Fig. S2), prompting us to examine how this surface might become accessible on RPA-coated ssDNA. Comparison of models of the RPA-ssDNA complex(*41*) with and without Polθ(*42*) revealed two alternative configurations. In the absence of Polθ, ssDNA engages the DNA-binding domains of both RPA1 and RPA2. In the Polθ-containing model, RPA2 DBD-D instead engages Polθ, while RPA1 retains contact with ssDNA through its DNA-binding domains (Fig. 4H, S8). These configurations suggest that Polθ docking may involve local disengagement of RPA2 from ssDNA without displacement of the entire RPA complex. We propose that such a configuration could be favored near the terminus of a resected DNA end, where limited ssDNA availability or transient disengagement of DBD-D may expose the Polθ-binding surface. Notably, the predicted complex positions RPA1 DBD-C and RPA2 DBD-D against Polθ, with ssDNA extending from RPA1 toward the DNA-binding channel of the Polθ helicase-like domain. This arrangement suggests that RPA may not only provide a docking platform for Polθ but also help position ssDNA for engagement by its helicase-like domain, potentially facilitating DNA handover. Together, these structural observations suggest that RPA may serve not merely as a docking platform for Polθ, but as a molecular scaffold that positions Polθ to engage the 3ʹ terminus of resected DNA.

## Discussion

DNA-end resection creates substrates for multiple DSB-repair pathways but the interactions that enable each pathway to use these substrates are poorly understood. Our findings identify an RPA2-Polθ interface essential for cellular TMEJ. Our new MUSIC-MAP technology identified a discrete RPA2 surface whose disruption selectively abolished TMEJ while preserving proliferation and HR, by multiple measures. Mutations in the complementary Polθ surface impaired TMEJ, whereas combining reciprocal charge-reversal mutations in RPA2 and Polθ restored TMEJ activity. These findings establish that cellular TMEJ depends on a specific interaction between RPA2 and Polθ, consistent with a role for RPA in enabling productive Polθ engagement at resected DNA ends.

This essential interaction challenges the prevailing view of RPA acting primarily as a barrier to TMEJ. RPA has generally been considered antagonistic to TMEJ because its binding to ssDNA suppresses annealing between short microhomologies at resected DNA ends(*31, 33*). Previous biochemical studies have shown that the Polθ helicase-like domain can displace RPA from ssDNA in an ATP-dependent manner(*43*). Our findings suggest that RPA recognition and displacement may represent sequential rather than opposing steps in TMEJ. Polθ may first engage RPA through the identified interface and subsequently remodel the RPA-ssDNA complex to facilitate microhomology search and annealing. Although the functional requirement for this interface is established here, the order and dynamics of these proposed steps remain to be determined.

The ability of TMEJ to act on resected DNA intermediates that are also substrates for HR raises the question of how Polθ accesses RPA-coated DNA ends. Structural modelling suggests how pathway specificity might be obtained. Local disengagement of RPA2 could expose the Polθ-binding surface while RPA1 retains contact with ssDNA, potentially allowing Polθ to engage a DNA terminus without displacing the entire RPA complex. The higher ssDNA-binding affinity of RPA1 DBD-A/B compared with DBD-C/D is consistent with a model in which DBD-C/D positions ssDNA for transfer to Polθ while DBD-A/B maintains its association with DNA(*44–46*). We propose that this configuration forms preferentially at ssDNA termini, positioning Polθ to guide the 3ʹ end into its helicase-domain channel(*42, 47*) and thereby facilitate microhomology annealing and subsequent DNA synthesis. In this model, RPA provides positional information that couples Polθ recruitment to productive engagement of the DNA end.

The RPA2-Polθ interaction may operate alongside mechanisms that regulate when and where Polθ is deployed. Recent studies have identified mitosis as an important context for TMEJ, in which replication-associated DSBs that persist into mitosis can undergo Polθ-dependent repair(*21*). TOPBP1-, RHINO-, and PLK1-dependent mechanisms promote Polθ activity at mitotic DNA lesions(*23, 24, 26*). These mechanisms may regulate Polθ deployment in a temporal and damage-context-dependent manner, whereas the RPA2 interaction could facilitate productive engagement with resected DNA ends. Layering these mechanisms would restrict an intrinsically mutagenic pathway to situations in which its capacity to eliminate persistent DNA breaks outweighs the mutations generated by repair.

The specificity of the RPA2 interface also exposes a potential therapeutic vulnerability. Polθ is being targeted in HR-deficient cancers because their survival can become dependent on TMEJ(*14, 15*). Existing inhibitors primarily target catalytic functions of Polθ. The finding that a single RPA2 substitution can essentially eliminate TMEJ without detectably compromising HR or proliferation demonstrates that pathway activity can instead be uncoupled through a protein-protein interaction. The corresponding Polθ surface therefore provides a conceptually distinct target for TMEJ inhibition that does not require inhibition of Polθ catalysis itself.

More broadly, our results point to a role for RPA beyond protecting ssDNA and coordinating its exchange between repair proteins. Human RPA was recently shown to interact directly with TERT and promote telomerase processivity(*48*). Notably, RPA2 W107, part of the Polθ-interacting surface identified here, is also critical for RPA-dependent stimulation of TERT. We therefore propose that RPA can function as an information-bearing platform: the organization of RPA on ssDNA exposes defined docking surfaces that license specialized enzymes at particular positions. Polθ engagement at resected DNA ends and TERT engagement at chromosome termini may thus represent examples of a broader mechanism by which RPA converts the geometry of ssDNA into spatially controlled enzymatic responses.

## Acknowledgments

We thank Sarah C. Moser for providing the HAP1-Cas9 cell line, and the LUMC protein facility and Susan Kloet and the staff of the Leiden Genome Technology Centre (LGTC) for technical support. The authors used ChatGPT, Claude and Copilot to assist with code generation, language editing, clarity, and organization of the manuscript text during preparation. The scientific content, analyses, interpretations, and conclusions were conceived, executed and verified entirely by the authors.

## Funding

This work was funded by grants from The Dutch Cancer Society (2021-2/13905 and 2024-3/16583) to M.T.

## Author contributions

Conceptualization: M.B., M.T.; Methodology: M.B.; Investigation: M.B., C.B., H.K., J.S., R.v.S.; Data interpretation: M.B., C.B., M.L., M.T.; Funding acquisition: M.B., M.T.; Supervision: M.B., M.T.; Writing: M.B., M.T.

## Competing interests

Authors declare that they have no competing interests.

## Data, code, and materials availability

Plasmids and cell lines generated are available upon request. Next-generation sequencing data will be made available upon publication of this manuscript. All other data are available in the main text or supplementary materials.

## Supplementary Materials

### Materials and Methods

#### Cell culture

129/Ola-derived IB10 mouse embryonic stem cell (mES cells) lines were seeded on 0.1% gelatin-coated dishes and cultured in complete medium (buffalo rat liver (BRL)-conditioned KnockOut DMEM (Gibco) supplemented with 100 U/mL penicillin, 100 μg/mL streptomycin, 2 mM GlutaMAX, 1 mM sodium pyruvate, 1× non-essential amino acids, 100 μM β-mercaptoethanol (all from Gibco), 10% fetal bovine serum (Gibco, #A5256701) and leukemia inhibitory factor, as previously described(*20*). The parental IB10-Cas9(*35*), *Polq^KO^*(*20*), *Lig4^KO^*(*20*) and *Trp53^KO^Palb2^KO^*(*49*) mES cell lines were generated previously. 293T cells were cultured in DMEM supplemented with GlutaMAX (both Gibco), 10% fetal bovine serum (Biowest, #S1600-500) and penicillin/streptomycin (Sigma), as previously described(*38*). HAP1 cells(*50, 51*) constitutively expressing Cas9 were generated and kindly gifted by Sarah C. Moser and were cultured in IMDM medium supplemented with 1% penicillin–streptomycin, 1% l-glutamine (all from Gibco) and 10% heat-inactivated fetal bovine serum (Biowest, #S1600-500). All cell lines were grown at 37 °C and 5% CO2 and were tested negative for mycoplasma contamination.

#### Plasmids

Plasmid pU6-(BbsI)_CBh-Cas9-T2A-mCherry was a gift from Ralf Kuehn (Addgene, #64324). pLenti-Cas9-2A-Blast was a gift from Jason Moffat (Addgene, #73310). pKLV2-U6gRNA5(BbsI)-PGKpuro2ABFP-W was a gift from Kosuke Yusa (Addgene, #67974). pLentiGuide-eGFP was a gift from Sylvie Noordermeer(*52*). Lentiviral packaging constructs pMDLg/pRRE, pRSV-Rev and pMD2.G were a gift from Didier Trono (Addgene, #12251, #12253, #12259).

SgRNA sequences were cloned into the pU6-(BbsI)_CBh-Cas9-T2A-mCherry backbone, or into the pKLV2-U6gRNA5(BbsI)-PGKpuro2ABFP-W backbone using two custom complementary DNA oligos (IDT) consisting of the target sequence and *BbsI* overhangs, which were melted, annealed and ligated (Promega, #M180B) into the corresponding *BbsI*-digested (NEB, #R3539S) plasmid backbone as described previously(*35, 54*). All constructs were sequence verified by Sanger. sgRNA sequences and corresponding oligos for cloning are provided in Table S1.

#### Transfections

Transfections of plasmids were performed using previously established methods(*38*). Briefly, mES cells were collected with trypsin, counted and resuspended in complete medium without pen/strep. Transfections were performed using Lipofectamine 2000 (Invitrogen, #11668019):DNA ratio of 2.4:1. Generally, separate mixtures containing 2 µg DNA in Opti-MEM (Gibco) and 4.8 µl Lipofectamine in 62.5 µl Opti-MEM (Gibco) were incubated for 5 min at room temperature (RT) before mixing and incubating for 20 min at room temperature. This mixture was added to the cells (1.0 × 10^6^ cells per transfection, resuspended in an equal volume), incubated in round-bottom tubes at 37 °C for 30 min, and subsequently seeded on gelatin-coated 12-well plates containing complete medium. The medium was refreshed the next day.

Transfections of RNP complexes were performed as previously described(*38*). Briefly, Alt-R CRISPR RNAs (crRNAs, ordered from IDT) were mixed with trans-activation CRISPR RNAs (tracrRNAs, ordered from IDT) at equimolar concentrations of 10 µM in nuclease-free duplex buffer (IDT) and incubated 5 min at 95 °C before gradually cooling down to RT to form RNA duplexes. Per 6-well transfection, RNP complexes were assembled by combining 37.5 µl RNA duplex (1 µM; diluted in nuclease-free duplex buffer) and 37.5 µl Cas9 protein (2 µM; produced by the LUMC protein facility) and diluted in Cas9 working buffer (20 mM HEPES; 150 mM KCl, pH 7.5)) in 550 µl Opti-MEM (Gibco), followed by incubation for 5 min at RT. Subsequently, RNP transfections were performed by adding 30 μL RNAiMAX (Invitrogen) and, when applicable for knock-in generation, 3.75 μL ultramer (10 μM in IDTE pH 7.5, both ordered from IDT) with 595 μL Opti-MEM followed by a 20 min incubation at RT before addition to 1.0 × 10^6^ mES cells or 7.5 × 10^5^ HAP1-Cas9 cells seeded in culture medium without antibiotics on 6-well plates. The medium was replaced with complete medium the following day to remove dead cells.

#### Lentivirus production

Lentivirus was produced in 293T cells by calcium phosphate transfection as previously described(*55*). Briefly, cells were seeded in 15-cm dishes one day before transfection and switched to antibiotic-free medium 2 hours before transfection. For each dish, 9 μg VSV-G envelope plasmid, 12.5 μg pMDLg/pRRE plasmid, 6.25 μg REV plasmid, and 32 μg transfer vector were mixed in 1125 μl 0.1× TE (pH 8.0), followed by the addition of 125 μl 2.5 M CaCl₂. After incubation for 5 min at RT, 1250 μl freshly prepared 2× HEPES-buffered saline (281 mM NaCl, 100 mM HEPES, 1.5 mM Na2HPO4 (Sigma-Aldrich, #71380, #H4034, #71643); adjusted to pH 7.12 with 5M NaOH) was added dropwise while vortexing. All solutions were 0.22-μm filter-sterilized before use. The resulting precipitate was immediately added to the cells, and the medium was replaced with complete medium appropriate for the target cell line the following morning. Virus-containing supernatants were collected 30 hours later, filtered through 0.45-μm PES syringe filters (Pall, #4614), aliquoted, and stored at −80 °C. Viral titers were determined by serial dilution and puromycin selection before use.

#### Generation of knockout and knock-in cell lines

Stable constitutive Cas9 expression in *Polq^KO^* or *Lig4^KO^* mES cells was established through lentiviral transduction with the pLenti-Cas9-2A-Blast plasmid, followed by multiple passages under blasticidin selection (5 µg/mL). *Rpa2*-hemizygous (*Rpa2 Δ/wild-type*) mES cells were generated by co-transfection of IB10-Cas9 cells with upstream (UPS) and downstream (DNS) targeting sgRNAs cloned into the pU6-(BbsI)_CBh-Cas9-T2A-mCherry backbone (Table S2). Single cells were seeded 48 hours after transfection and individual colonies were picked 7 days later. Clones with the hemizygous genotype were confirmed using PCR primers to amplify the upstream target-region, the downstream target-region, the delta-allele, and the wild-type allele followed by Sanger sequencing (Table S3). *Polq*-hemizygous (*Polq Δ/wild-type*) mES cells were generated and verified using a similar strategy.

Monoclonal knock-in cell lines were generated by RNP transfection of *Rpa2 Δ/wild-type* mES cells with a crRNA (IDT) targeting the vicinity of the knock-in site (Table S4) whilst supplying a variant-specific ultramer donor (IDT) to edit the remaining wild-type allele into the desired variant allele (Table S1). Single cells were seeded 48 hours after transfection and single colonies were picked 7 days later. Clones were verified by PCR and Sanger sequencing (Table S3). The editing of *Polq* in *Rpa2 Δ/D109R* mES cells was performed in the presence of 1 µM DNA-PKi (AZD7648, MedChemExpress, #HY-111783) for 2 days to improve knock-in efficiency.

Monoclonal *RPA2 D109R* and *RPA2 W107A-D109A* knock-in HAP1-cas9 cell lines were generated by RNP transfection in the presence of 2 µM DNA-PKi (AZD7648, MedChemExpress, #HY-111783) and 10 µM Polθ inhibitor (ART558, kindly gifted by Artios Pharma) for 2 days to improve knock-in efficiency.

#### Rpa2-focused sgRNA library construction

To scan (nearly) all sgRNAs targeting the exons of the *Rpa2* gene for their effect on DNA repair mutational outcome distributions, sgRNA sequences were extracted using InDelphi software (*36*) and a custom library was constructed as described previously(*35*). Corresponding sgRNA sequences, to which overhangs were added for cloning, were ordered as oPools (IDT) (Data S1) and cloned into the pKLV2-U6gRNA5(BbsI)-PGKpuro2ABFP-W backbone using the NEBuilder HiFi DNA assembly Master Mix (NEB, #E2621L) according to the manufacturer’s instructions. Assembly reactions contained 50 ng BbsI-HF digested (NEB, #R3539S) and gel-purified (Qiagen, #28704) plasmid, 5 µl of diluted oPool (0.2 µM in 1× NEBuffer 2.0), 2× NEBuilder Hifi Assembly Master Mix, and nuclease-free water to a final volume of 20 µl. Assembly reactions were incubated for 1 hour at 50 °C and purified using the QIAquick PCR Purification Kit (Qiagen; Catalogue# 28106) following manufacturers’ protocol. Next, assembled products were electroporated into Endura Duo electrocompetent cells (Lucigen, #60242-1) using a MicroPulser electroporator (Bio-Rad; Ec1 programme) and recovered in ROC Expression Recovery Medium (Lucigen, #80030-1). Library representation was determined by serial dilution plating on ampicillin-containing agar plates and exceeded 10,000-fold per sgRNA. The remaining transformed culture was expanded in ampicillin-containing 2x LB medium at 37 °C overnight, harvested, and plasmid libraries were recovered using a NucleoBond-based purification procedure (Macherey-Nagel, #740410.50). A control library containing 5 sgRNAs targeting mm*Lig4*, 5 sgRNAs targeting mm*Polq*, and 30 non-targeting sgRNAs (extracted from (*56*)) was generated using the same procedure and yielded an equivalent library representation after cloning.

#### Mutational scars of induced DNA cleavage (MUSIC) scanning of Rpa2

IB10-Cas9 mES cells were transduced in bulk with the focused mm*Rpa2* and control-library (mixed in a 1:3 ratio during lentiviral production). A total of 4.0 × 10^7^ cells were transduced at an MOI of ∼0.2 in the presence of 8 µg/mL polybrene. The next day, cells were placed under puromycin selection (2 µg/mL) for 3 days, during which the cells were passaged once. Cells were subsequently cultured in complete medium without puromycin, and repair outcome profiles were generated 5 days after transduction. To this end, cells were transduced a second round, but now using lentivirus to express the sg*Targeting-1* sgRNA from the pLentiGuide-eGFP plasmid at an MOI of 2-5. This sgRNA targets a unique region neighboring the sgRNA expression cassette within the pKLV2-U6gRNA5(BbsI)-PGKpuro2ABFP-W backbone, enabling repair outcomes to be linked to their corresponding genetic backgrounds by short-read sequencing(*35*). Culture medium was replaced the following day, and cells were collected and aliquoted 3 days after transduction, ensuring adequate cell numbers to preserve library representation. Replicates were derived from a single bulk introduction of the sgRNA library, split into 3 populations before the second round of lentiviral targeting, when repair profiles are generated, and maintained separately throughout all subsequent steps.

#### MUSIC based Mapping of Allelic variants to Protein function (MUSIC-MAP)

To map allelic variants to protein function, the MUSIC workflow was slightly adapted.

For the MUSIC-MAP of *Rpa2* in mES cells, an ssODN template library was designed to cover the residues predicted by AlphaFold to mediate the Polθ-interaction (AA105-109). The library comprised single substitutions of each residue to either a neutral residue (alanine) or a disruptive residue (R-to-E, D-to-R, W-to-S). Combinatorial double, triple and quadruple substitution allele variants were included to assess their combined effects, alongside the two predicted 9 bp deletion alleles naturally generated by the *Rpa2*-targeting sgRNA identified in the MUSIC library for validation. For each allele variant, two synonymous sequences were designed as internal replicates. Six synonymous wild-type sequences were used to derive reference mutational outcome spectra. Each variant sequence was flanked by ∼65 bp homology arms complementary to the target locus to facilitate single-strand template repair. The ssODN template library (Data S1) was ordered as oPool (IDT) and supplied during RNP transfection of IB10-Cas9 *Rpa2 Δ/wild-type* cells, generating a pooled population in which the remaining wild-type allele harbored individual engineered variants. Transfections were performed on 9.0 × 10^6^ mES cells per replicate, generating variant populations that were estimated, on the basis of transfection efficiency, to contain thousands of independently derived knock-in clones per template. Variant representation was monitored and maintained throughout the experiment by cell counting (Beckman Z2 Coulter Particle Count and Size Analyzer). Cells were expanded for 7 days before lentiviral targeting (1.7 × 10^8^ cells per replicate distributed over 16× 15-cm plates, MOI 2-5, 8 µg/mL polybrene) of the adjacent intron (Table S2), yielding complex mutational spectra comprising tens to hundreds of thousands of individual repair products. Culture medium was replaced the next morning for complete medium containing 2 µg/mL puromycin, and refreshed a second time on day 2. In parallel, a single 15-cm plate was transduced in the presence of 10 µM ART558 (kindly gifted by Artios pharma) and used as a reference for a Polθ-deficient repair profile. Cells were harvested and aliquoted at sufficient representation to maintain library complexity 30 hours after targeting. Replicates consisted of two independent RNP transfections that were maintained as separate cell populations throughout all subsequent steps.

For the MUSIC-MAP analysis of *POLQ* in HAP1-Cas9 cells, an ssODN template library was designed to target residues on the reciprocal Polθ interaction surface (E691, R696, K699, and K701; Data S1). Given the smaller size of the library (47 variants), 4 synonymous wild-type sequences were included to derive control repair spectra. Screening of the library was performed using the RNP procedure described above, but scaled to HAP1-Cas9 format (4.5 × 10^6^ cells per bulk transfection, Table S4). After 7 days of culture, the adjacent intron was targeted using lentivirus to express the corresponding sgRNA (Table S2). Since the crRNA targets a *POLQ*-exon, Cas9-induced frameshift alleles served as the internal reference for a Polθ-deficient repair profile. Replicates were derived from a single bulk RNP transfection to generate a pooled population of genetic backgrounds, split into two independent populations before lentiviral targeting for repair outcome profiling and maintained separately throughout all subsequent steps.

For the MUSIC-MAP of *Polq* in mES cells, the ssODN template library consisted of two synonymous wild-type variants and further included the single Polθ R693D, Polθ K698D and combinatorial Polθ R693A-K698A, Polθ R693A-K698D, Polθ R693D-K698A and Polθ R693D-K698D variants (Data S1). This library was screened in both IB10-Cas9 *Polq Δ/wild-type* and IB10-Cas9 *Rpa2 Δ/D109R* mES cells using RNP transfections (1.0 × 10^6^ cells per transfection, Table S4) with the inclusion of 1 µM DNA-PK inhibitor (AZD7648; MedChemExpress, #HY-111783) to generate pooled variant populations. DNA-PK inhibitor was washed out of the culture medium 2 days after targeting and repair spectra were generated on day 10 by lentiviral transduction (4.0 × 10^7^ cells, MOI 5) to express an sgRNA targeting the adjacent intron (Table S2). Replicates consisted of three independent RNP transfections that were maintained as separate cell populations throughout all subsequent steps.

#### Repair outcome profiling in monoclonal cell lines

Mutational profiles of non-multiplexed samples were derived by lentiviral targeting of the indicated genomic site to express the corresponding sgRNA from the pLentiGuide-eGFP backbone (MOI 2-5), unless otherwise indicated. Per transduction, monoclonal mES cells (5.0 × 10^5^) or HAP1-Cas9 (3.0 × 10^5^) were seeded on 0.1% gelatin-coated (only for mES cells) 6-well plates in complete medium containing 8 µg/mL polybrene. Cells were selected with puromycin (2 µg/mL for mES cells, 1 µg/ml for HAP1-Cas9) the next day and harvested 72 hours after transduction. The *HBB* locus was targeted in HAP1-Cas9 cells (1.5 × 10^5^) using RNP transfection and cells were harvested 4 days later. Samples were stored as frozen cell pellets at −20 °C. The initial representation was maintained throughout all steps of culturing and subsequent library preparation.

#### Genomic DNA isolation from cell lines

Genomic DNA was isolated from screening samples using the Blood & Cell Culture DNA Kit (Qiagen, #13362, #10262, and #19060) according to the manufacturer’s instructions for cell culture samples and scaled to the recommended loading capacity per Genomic-tip.

Genomic DNA isolation from frozen pellets of monoclonal cell lines was performed using established methods(*38*), unless stated otherwise. Cells were lysed using 10mM Tris-HCL pH7.5, 10mM EDTA, 200mM NaCl, 1% SDS and 0.4mg/mL Proteinase K (up to 4e^6^ cells per 500 μL volume) and incubated at 55°C for 3-16 hours. Next, 0.3x volume saturated NaCl (0.36 g/mL) was added, and proteins were precipitated by centrifugation at 20,000g for 10 min. DNA was collected by transferring the supernatant to a clean tube containing 0.7x of the total volume isopropanol, followed by centrifugation and washing of the pellet with 70% ethanol before resuspension in TE buffer (pH 8.0).

#### Genotyping of monoclonal cell lines

All knock-in and knockout cell lines generated in this study were genotyped by PCR on the target loci using GoTaq (Promega, #M7848) according to the manufacturer’s instructions with the following PCR program: (1) 95 °C for 10 min, (2) 35 cycles of 95 °C for 30 s, 60 °C for 30 s, 72 °C for 60 s, (3) 72 °C for 5 min. PCR amplification was performed using locus-specific primers under the conditions described below. Amplified products were analysed by gel electrophoresis, diluted 5× in MQ and 2 µl of diluted product was submitted for Sanger sequencing. Primer sequences and expected amplicon sizes for all assays are provided in Tables S1 and S3.

#### NGS library preparation for Cas9-induced repair outcome profiling

All sample preparations were scaled to maintain library representation, using 6 µg and 7.2 µg gDNA per million mouse and human genomes, respectively.

Sample preparation of samples in which MUSIC was applied to an *Rpa2*-focused library was performed as previously described(*35*). The target region was PCR amplified using NEBNext Ultra II Q5 Master Mix (NEB, #M0544) with target-site specific primers (IDT) that contain Illumina overhang adaptors (Table S1) and the following PCR program: (1) 98 °C for 3 min, (2) 18 cycles at 98 °C for 10 s, 62 °C for 30 s, 72 °C for 45 s, (3) 72 °C for 5 min. An input of 4 µg genomic DNA and 0.2 µM oligo was used per 50 µl parallel reaction during this first round of PCR. All parallel reactions of the same sample were then pooled and an excess was purified using AMPure XP Beads (Beckman Coulter, #A63882) in a 0.8× ratio and eluted in MQ. The second PCR incorporated unique P5 and P7 sequences and Illumina flow-cell adaptor sequences in a single-step through nested primers using NEBNext Ultra II Q5 Master Mix (NEB, #M0544) containing 5 µl of purified PCR1 template and 0.2 µM oligo (IDT) with the following PCR program: (1) 98 °C for 2 min, (2) 12 cycles at 98 °C for 10 s, 65 °C for 75 s, (3) 65 °C for 5 min. The reaction mixture was purified using magnetic AMPure XP Beads in a 1.0× ratio and eluted in 25 µl MQ. The final PCR amplicon (378 bp, excluding adaptors) contained the DSB target site and sgRNA sequence 75 bp and 35-55 bp from the forward and reverse primers, respectively.

MUSIC-MAP samples were processed similarly, with the following changes: Illumina flow-cell adaptor sequences were added in the first PCR with the following program: (1) 98 °C for 3 min, (2) 18 cycles at 98 °C for 10 s, 60 °C for 30 s, 72 °C for 45 s, (3) 72 °C for 5 min. Parallel reactions were purified using AMPure XP Beads in ratios of 1.2×, 1.2× and 1.0× for mouse *Rpa2*, human *POLQ* and mouse *Polq*, respectively. The second PCR incorporated unique P5 and P7 sequences using universal primers (Table S1), purified in ratios of 1.2×, 1.2× and 0.8× for mouse *Rpa2*, human *POLQ* and mouse *Polq*, respectively. For mouse *Rpa2* in hemizygous-*Rpa2* mES cells, the final PCR amplicon (280 bp, excluding adaptors) contained allele variant sequences 102-123 and the DSB target site at 81 bp from the forward and reverse primers, respectively. For human *POLQ* in HAP1-Cas9 cells, the final PCR amplicon (275 bp, excluding adaptors) contained the DSB target site at 73 bp and the allele variant at position 112-145 bp from the forward and reverse primer, respectively. For MUSIC-MAP of *Polq* in mES cells, the final PCR amplicon (481 bp product, excluding adaptors) contained the allele variant at position 123-142 bp and the DSB target site at 96 bp from the forward and reverse primer, respectively.

NGS library preparation from samples in which monoclonal cell lines were targeted were performed using the same approach as for MUSIC-MAP, but scaled to ≥2.5 × 10^5^ cells in a single 50 µl reaction and using locus-specific primers (Table S1). The second PCR was scaled to a single 25 µl reaction. Amplicons were purified using AMPure XP Beads in ratios of 1.2× and 1.0× following PCR1 and PCR2, respectively.

DNA quantity and quality were monitored throughout library preparation using a NanoDrop spectrophotometer (Thermo Fisher Scientific, ND-1000), a Qubit 2.0 fluorometer (Invitrogen; Catalogue# Q32866) with the dsDNA High Sensitivity Assay Kit (Invitrogen; Catalogue# Q32854), and agarose gel electrophoresis. Final library size distribution and purity were assessed using a High Sensitivity DNA Kit (Agilent; Cat. No. 5067-4626) on a 2100 Bioanalyzer (Agilent). Libraries were pooled at equimolar ratios and sequenced as 150-bp paired-end reads on a NovaSeq X platform (Illumina).

#### Bioinformatic analysis of outcome spectra

Repair outcome spectra were derived from paired-end reads as previously described (*34, 35*). For pooled samples, sequencing reads were first demultiplexed based on exact matching of expected sgRNA (MUSIC) or variant-allele (MUSIC-MAP) sequences present between predefined left and right flanking sequences. Specifically: for MUSIC samples the sgRNA sequences were identified in the R2 read and the corresponding R1 reads were extracted and binned.

For MUSIC-MAP of *Rpa2* in mES cells, wild-type and predicted 9 bp deletion alleles were included in the expected sequences during demultiplexing as internal controls for the repair profiles of their corresponding synonymous ssODN-engineered variants. Paired-end reads (R1 and R2) were merged prior to demultiplexing and trimmed before subsequent SIQ analysis on binned demultiplexed samples. With the exception of analyses comparing crRNA-induced and ssODN-induced variants, all subsequent analyses were confined to ssODN-induced alleles.

For MUSIC-MAP of *POLQ* in HAP1-Cas9, the most common crRNA-induced frameshift alleles were included in the expected sequences to serve as internally controlled reference profiles of genetic Polθ depletion. Paired-end reads (R1 and R2) were merged prior to demultiplexing and trimmed before binning and subsequent SIQ analysis. During our analysis, SIQ identified events that were exclusive to specific ssODNs and observed in both replicates, but never in the ssODN designed to introduce the same alteration. We attribute these events to errors introduced during ssODN synthesis, which are copied into the genome through HDR. Events that accounted for >1% of the total mutational spectrum, were unique to a single ssODN and were observed in both replicates were excluded from further analysis.

For MUSIC-MAP of *Polq* in *Rpa2* wild-type and D109R mES cells, the most common crRNA-induced frameshift alleles were included in the expected sequences to serve as internally controlled reference profiles of genetic Polθ depletion. Variant alleles were identified in the R1 read and the corresponding R2 reads were extracted and binned.

The demultiplexed reads were binned and analyzed using SIQ software (version 2.2a) to assign repair outcomes from sequencing reads and the output was analyzed and plotted using the corresponding SIQplotteR web tool, as previously described (*34, 35*). Samples in which repair spectra were generated in monoclonal cell lines were directly analyzed using SIQ software as above. SIQ classifies reads into the following events: deletion, deletion with an insertion of de novo DNA at the location of the deletion (delins), insertion (separated into 1 bp or >1 bp length), templated insertion (≥6 bp insertion that can reliably be mapped to the flanking sequence(*34*)), HDR (only for MUSIC, a specific deletion-insertion event that can be mapped to the viral backbone used in the second round of targeting), SNV (single-nucleotide variant) or wild-type (identical to the reference sequence). Additional parameters, including deletion size and microhomology length (for deletions) are extracted to allow further grouping. Outcomes were only included in the final data if at least two reads supported that event, and analyses were restricted to non-SNV outcomes mapped within a vicinity of ≤2 bp of the predicted cleavage site. Relative mutagenic outcome fractions were calculated using only non-wild-type reads.

From these data, the log2 fold-change of each outcome was calculated per sgRNA (MUSIC) or variant allele (MUSIC-MAP) relative to controls (non-targeting sgRNAs or sequences matching synonymous wild-type encodings, respectively). Significance was assessed using a binomial test (binom.test, R Stats Package) with the mean outcome frequency across control samples used as the expected probability of success. For each sgRNA or variant allele, the number of reads corresponding to each outcome was used as the number of successes, and the total number of mutagenic reads as the number of trials. P values were transformed as −log10 and are reported as P-score. To account for multiple testing, the significance threshold was scaled to the number of sgRNAs or variant alleles in the library, applying a Bonferroni-adjusted cutoff of P-score ≥ 3 (adjusted P = 0.001). Heatmaps were generated using the heatmap.2 function from the gplots package, including the 100 most abundant outcomes whose frequencies were significantly altered by at least one sgRNA (MUSIC) or variant allele (MUSIC-MAP) in all replicates. UMAP embeddings were subsequently generated from the heatmap data using the umap package.

Tornado plots were generated using the SIQplotteR web tool. Where indicated, samples were pooled within SIQplotteR by assigning equal weight to each sample irrespective of read count. Outcome allele and outcome-type (templated insertion) frequencies were extracted from the corresponding repair profiles. Statistical testing and data visualization were performed in GraphPad Prism (v10.6.1).

#### Cell proliferation assay

For growth curve analyses, cells were trypsinized, counted, and seeded in technical duplicates at 1.0 × 10³ cells per well in 0.1% gelatin-coated 96-well plates. The numbers of viable cells were quantified at 24 hour intervals using the CellTiter-Glo Luminescent Cell Viability Assay (Promega, #G7571). CellTiter-Glo reagent was added at a 1:1 ratio to the culture medium, followed by orbital shaking at 1,000 rpm for 2 min and incubation at room temperature for 10 min. Luminescence was measured in black-well/clear-bottom plates using the SpectraMax iD3 plate reader (Molecular Devices). Background-subtracted luminescence values from technical duplicates were averaged and normalized to the initial time point.

#### Cell cycle analysis

To obtain cell cycle profiles from mES cells, cells were harvested, counted, and 5.0 × 10^5^ cells were washed in ice-cold PBS followed by overnight fixation in ice-cold 70% ethanol in 1x PBS at 4 °C. Prior to flow cytometry using a Novocyte Flow Cytometer (ACEA Biosciences, Inc.), cells were pelleted, washed in PBS containing 2% FCS, and subsequently resuspended in a fixed amount of PBS containing 2% FCS, 25 μg/ml propidium iodide (Sigma, #P4170), 100 μg/ml RNase A (Sigma, #R6513) and incubated at 37 °C for 30 min. Stained cells were transferred to FACS tubes and analyzed immediately. Cell cycle profiles were obtained from viable singlet cells, identified by gating, and analyzed using the Watson (Pragmatic) model in the instrument software, with unconstrained G1 peak, G1 CV, G2 peak and G2 CV parameters.

#### IR-survival assay

The sensitivity to ionizing radiation was determined by seeding 0.25-10 × 10^3^ mES cells in 0.1% gelatin-coated 6-cm plates in technical duplicates per condition. Cells were treated with 0-6 Gy of radiation using the YXlon Maxishot X-ray generator (200 KV, 12 mA, dose rate ∼4 Gy/min) irradiator 3 hours after plating. Combination treatment of IR with the Polθ inhibitor ART558 was performed by adding 10 μM ART558 during plating. Seven days later, plates were washed with 0.9% NaCl, stained with methylene blue (Sigma, #M9140) and colonies (≥50 cells) were counted manually using an inverted microscope. Colony counts from technical duplicates were combined, and experiments were performed in three independent biological replicates. Data were fitted to a linear-quadratic model in GraphPad Prism (v10.6.1). Statistical significance was assessed using the CFAssay software package (v1.44.0)(*57*), which performs ANOVA-based testing, and resulting P values were corrected for multiple testing using the Benjamini–Hochberg procedure in R.

#### PARPi sensitivity assay

For proliferation-based PARPi sensitivity assays, mES cells were seeded in triplicate at 1.0 × 10^4^ cells per well onto 0.1% gelatin-coated 96-well plates, with the indicated concentration of KU-0058948 hydrochloride (MedChemExpress, #HY-136489A). After 96 hours, cells were collected and quantified using a Novocyte Flow Cytometer (ACEA Biosciences). Viable singlet cells were identified by gating and cell counts from technical triplicates were averaged and normalized to the untreated control.

#### Immunofluorescence

IR induced 53BP1- and RAD51 foci formation on mES cells was performed as previously described(*58*), with minor modifications. Exponentially growing cells were exposed to 5 Gy of radiation using the YXlon Maxishot X-ray generator (200 KV, 4 mA, dose rate ∼1.2 Gy/min) or left untreated. One hour after IR, cells were collected with trypsin and seeded (1.25 × 10^5^ and 2.0 × 10^5^) cells per chamber as technical duplicate) onto 8-well chamber slides precoated with Poly-L-Lysine (Merck, #A-005-C). Three hours after IR, cells were washed once with PBS++ (PBS supplemented with 1mM CaCl_2_ and 0.5mM MgCl_2_), followed by pre-extraction using ice-cold CSK buffer (10 mM HEPES KOH pH 7.8, 100 mM NaCl, 300 mM Sucrose, 3 mM MgCl_2,_ 1 mM EGTA and 0.5% v/v Triton X100) on ice for 5 min and ice-cold CSS buffer (10 mM Tris pH 7.5, 10 mM NaCl, 3 mM MgCl_2,_ 1% v/v Tween and 0.5% w/v Sodium Deoxycholate) on ice for 5 min. Cells were washed with PBS++ and fixed with 2% PFA in PBS++ for 20 min at RT. PFA was removed and cells were washed 3 times with PBS++ and stored at 4 °C. Prior to staining, cells were permeabilized with 0.2% Triton X-100 in PBS++ for 20 min. Cells were washed 3 times in staining buffer (PBS++, BSA (1%), glycine (0.15%), Triton X-100 (0.1%)) and then incubated for 30 min in staining buffer at RT. Slides were incubated with primary antibodies (rabbit anti-53BP1, Novus, #NB100-304, 1:5000; and rabbit anti-RAD51, BioAcademia, #70-001 lot.04, 1:20.000 in staining buffer) for 1 hour before washing 3 times with staining buffer for 5 min each. The secondary antibody (Alexa Fluor 555 goat anti-rabbit, Invitrogen, #A21429 lot:2790124, 1:1000 in staining buffer), supplemented with DAPI and incubated for 1 hour. Cells were washed three times in staining buffer, once in PBS++, and then mounted in Aqua-Poly/Mount (Polysciences, #18606) before imaging on a Leica SP8 upright confocal microscope equipped with an HC PL APO 40×/1.30 oil-immersion objective. Foci counts per nucleus were quantified using the FociAnalyzer plug-in in Fiji(*59*). For each replicate and condition, at least four fields were imaged, quantified and averaged to obtain a replicate mean. The experiment was performed at least three times and statistics were calculated on replicate means.

#### AlphaFold structure prediction and complex modeling

Structural models of mouse RPA–Polθ complexes were generated using AlphaFold 3(*37*). Multiple independent predictions were performed using default parameters, with structural templates excluded. Models were ranked using predicted local distance difference test (pLDDT) and interface predicted template modeling (ipTM) scores. Initial predictions included *Mus musculus* (mm) mmRPA2 DBD-D (residues 48–178), mRPA1 DBD-C (residues 454–623), and the mmPolθ helicase domain (residues 63–889), together with a 14-nucleotide poly(dT) ssDNA strand, yielding a model with ipTM = 0.77 and pTM = 0.83. Subsequent predictions included mmRPA2 (residues 48–178), mmRPA1 (residues 207–623), mmRPA3, and mmPolθ (residues 43–889), together with a 30-nucleotide poly(dT) ssDNA strand, yielding a model with ipTM = 0.70 and pTM = 0.73. For comparison, the corresponding complex was modeled without Polθ, yielding ipTM = 0.74 and pTM = 0.63. To model complexes containing charge-reversal substitutions, predictions were performed using the same RPA–Polθ complex and 30-nucleotide poly(dT) strand, with RPA2-D109R combined with Polθ-K698D (ipTM = 0.72, pTM = 0.76). Predicted complexes were visualized and analyzed using UCSF ChimeraX(*60*). Potential contacts at the RPA2– Polθ interface were identified by manual inspection using a distance cutoff of <3 Å. The predicted RPA1–RPA2–RPA3–Polθ–ssDNA complex was superposed onto the experimentally determined *Homo sapiens* DNA-bound Polθ helicase-domain dimer (PDB 9C5Q(*42*)), using Polθ as the alignment reference, and onto the *Pyrococcus abyssi* RPA–ssDNA filament structures (PDB 8OEJ and 8OEL(*41*)), using RPA2 as the alignment reference. All structural figures were prepared using UCSF ChimeraX.

#### Statistics and software versions

Mutational spectra were generated using SIQ (version 2.2a). Corresponding plots were made using the packages ggplot2 (version 3.5.2) and umap (version 0.2.10.0) in RStudio (version 2025.05.0) and the SIQplotteR web tool. GraphPad Prism (version 10.6.1) was used for visualization and statistical analysis of bar graphs and survival curves. Statistical significance was calculated as indicated in each figure legend. Analyses of IR-survival curves were performed using the CFAssay package (version 1.44.0) in R (version 4.5.1).

### Supplementary Figures

**Figure S1.**
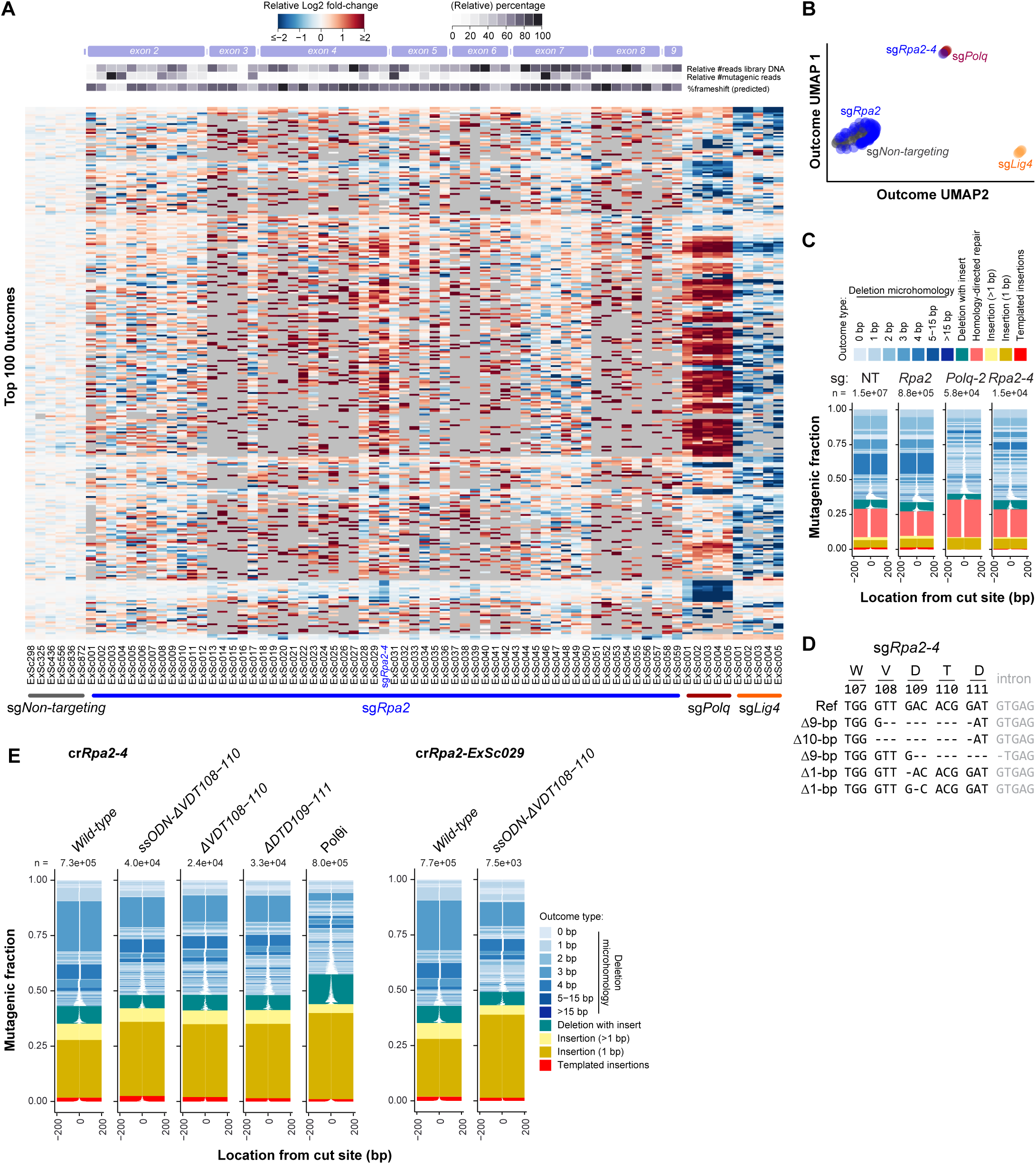
Identification and validation of RPA2 alleles that selectively impair TMEJ. (**A**) Heatmap of a MUSIC-based exon scan of *Rpa2* in Cas9-expressing mES cells. A custom sgRNA library targeting *Rpa2* exons was screened in bulk using MUSIC. The heatmap shows the log2 fold-change in frequency of the top 100 mutagenic repair outcomes (y-axis, grouped by outcome type) for each sgRNA (x-axis, grouped by target gene). Three biological replicates, representing independent transductions of a pooled cell population, are shown individually. Six non-targeting sgRNAs are included for comparison. The positions of *Rpa2*-targeting sgRNAs relative to the targeted exons are indicated schematically above the heatmap (not to scale). Relative sgRNA abundance was determined by sequencing the library DNA after cloning and at the experimental endpoint, and is shown as a percentage of the most abundant sgRNA. Predicted frameshift frequencies were calculated using inDelphi(*36*). Grey indicates outcomes for which no supporting reads were detected. (**B**) UMAP projection of the repair profiles shown in (A), revealing clustering of *sgRpa2-4* with *Polq*-targeting sgRNAs. (**C**) Tornado plots showing the complete spectrum of mutagenic repair outcomes. Individual repair products are plotted according to their frequency among mutagenic reads (y-axis) and position relative to the cleavage site (x-axis). Outcomes are colored by repair product type, with white regions indicating deleted sequences. sgNT and sg*Rpa2* represent pooled repair profiles from non-targeting and *Rpa2*-targeting sgRNAs, respectively, whereas *sgPolq-2* and *sgRpa2-4* represent individual sgRNAs. Biological replicates were combined for visualization. The total number of mutagenic reads is indicated above each plot. bp, base pairs. (**D**) In silico prediction of the predominant editing outcomes generated by *sgRpa2-4*, including two distinct in-frame deletions. (**E**) Tornado plots showing the complete spectrum of mutagenic repair outcomes in cells carrying defined 9 bp in-frame deletions at the endogenous *Rpa2* locus. The ssODN-ΔVDT108–110 profiles were obtained from cells in which the ΔVDT108–110 deletion was introduced using ssODN donors containing additional synonymous substitutions and two independent crRNAs (cr*Rpa2-4* and cr*Rpa2-ExSc029*) to control for potential off-target effects. For comparison, repair profiles of cells carrying 9 bp in-frame deletions generated through microhomology-mediated repair following cleavage with cr*Rpa2-4* are also shown. Treatment with the Polθ inhibitor ART558 (10 µM) served as a TMEJ-deficient control. Individual repair products are plotted according to their frequency among mutagenic reads (y-axis) and position relative to the cleavage site (x-axis). Outcomes are colored by repair product type, with white regions indicating deleted sequences. The total number of mutagenic reads is indicated above each plot.

**Figure S2.**
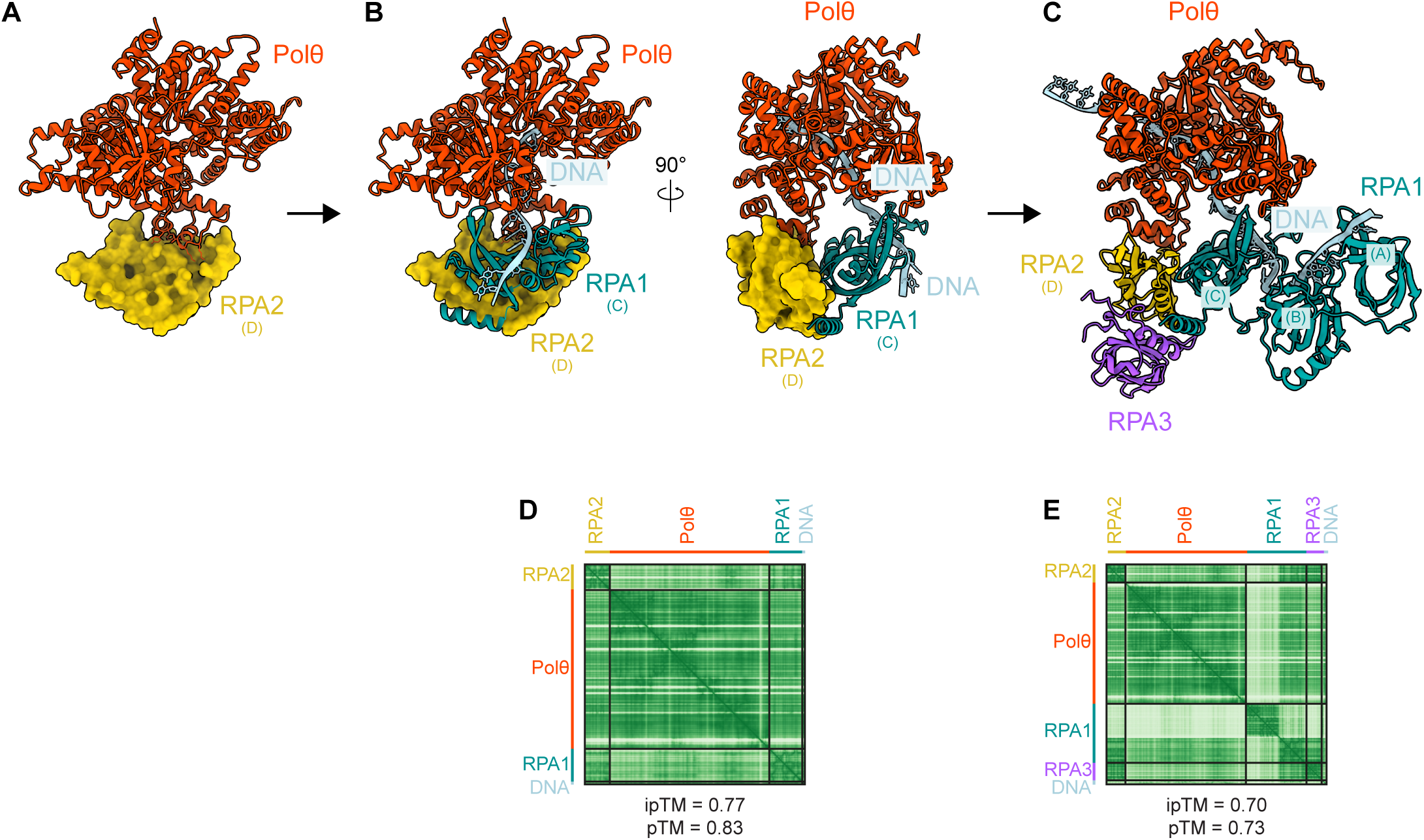
AlphaFold3 modeling predicts an interaction between the RPA2 ssDNA-binding domain D and the N-terminal helicase domain of Polθ. (**A–E**) AlphaFold3 was used to model the interaction of Polθ (mouse) with RPA (mouse) bound to ssDNA. An initial model incorporating the Polθ N-terminal helicase domain, the OB-folds of RPA1 (domain C) and RPA2 (domain D), and a 14-nt ssDNA substrate predicted an interface between Polθ and RPA2, involving RPA2 residues 106–109, with additional contacts involving RPA1 domain C (**A,B**). The model showed high confidence scores (pTM = 0.83, ipTM = 0.77), with the predicted aligned error (PAE) plot supporting the relative positioning of the interacting domains (**D**). To assess whether this interface is compatible with the larger RPA complex, the Polθ helicase domain was modeled together with RPA1 domains A–C, RPA2 domain D, RPA3, and a 30-nt ssDNA substrate. This expanded model retained the predicted Pol-RPA2 interface (**C**) and showed similarly high confidence scores (pTM = 0.73, ipTM = 0.70; **E**).

**Figure S3.**
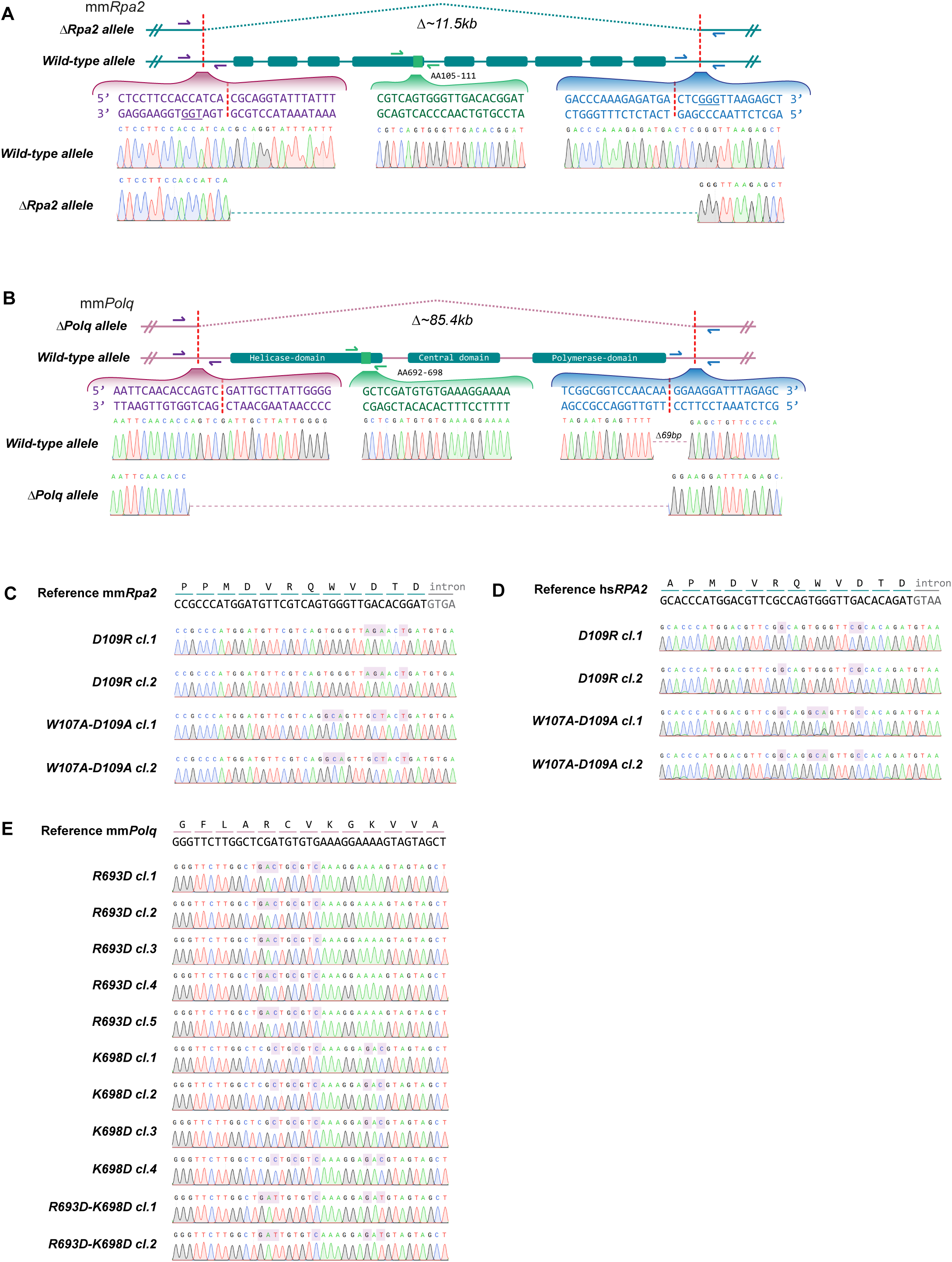
Generation and validation of clonal RPA2 and Polθ mutant cell lines. (**A,B**) Schematic representation and Sanger sequencing validation of monoclonal mES cell lines hemizygous for *Rpa2* (**A**) and *Polq* (**B**). The *Rpa2* deletion spans approximately 11.5 kilobases (kb), whereas the *Polq* deletion spans approximately 85.4 kb. The targeted regions and corresponding wild-type and deletion alleles are shown, together with representative sequencing chromatograms confirming the genomic deletions. Of the remaining wild-type alleles, wild-type sequence is retained around the crRNA induced cut sites, except for the downstream region of *Polq*, which showed a deletion of 69 base pairs. (**C,D**) Sanger sequencing validation of independently derived mES (*Rpa2* hemizygous background) (**C**) and human *HAP1* (**D**) clones carrying the indicated RPA2 interface mutations (D109R or W107A-D109A). Reference sequences and corresponding amino acid positions are shown above the sequencing chromatograms. Differentiating nucleotides are marked by light magenta squares. (**E**) Sanger sequencing validation of independently derived mES clones carrying the indicated *Polq* substitutions, individually or in combination, in the cellular background of RPA2 Δ/D109R. Reference nucleotide and amino acid sequences are shown above the sequencing chromatograms. Differentiating nucleotides are marked by light magenta squares.

**Figure S4.**
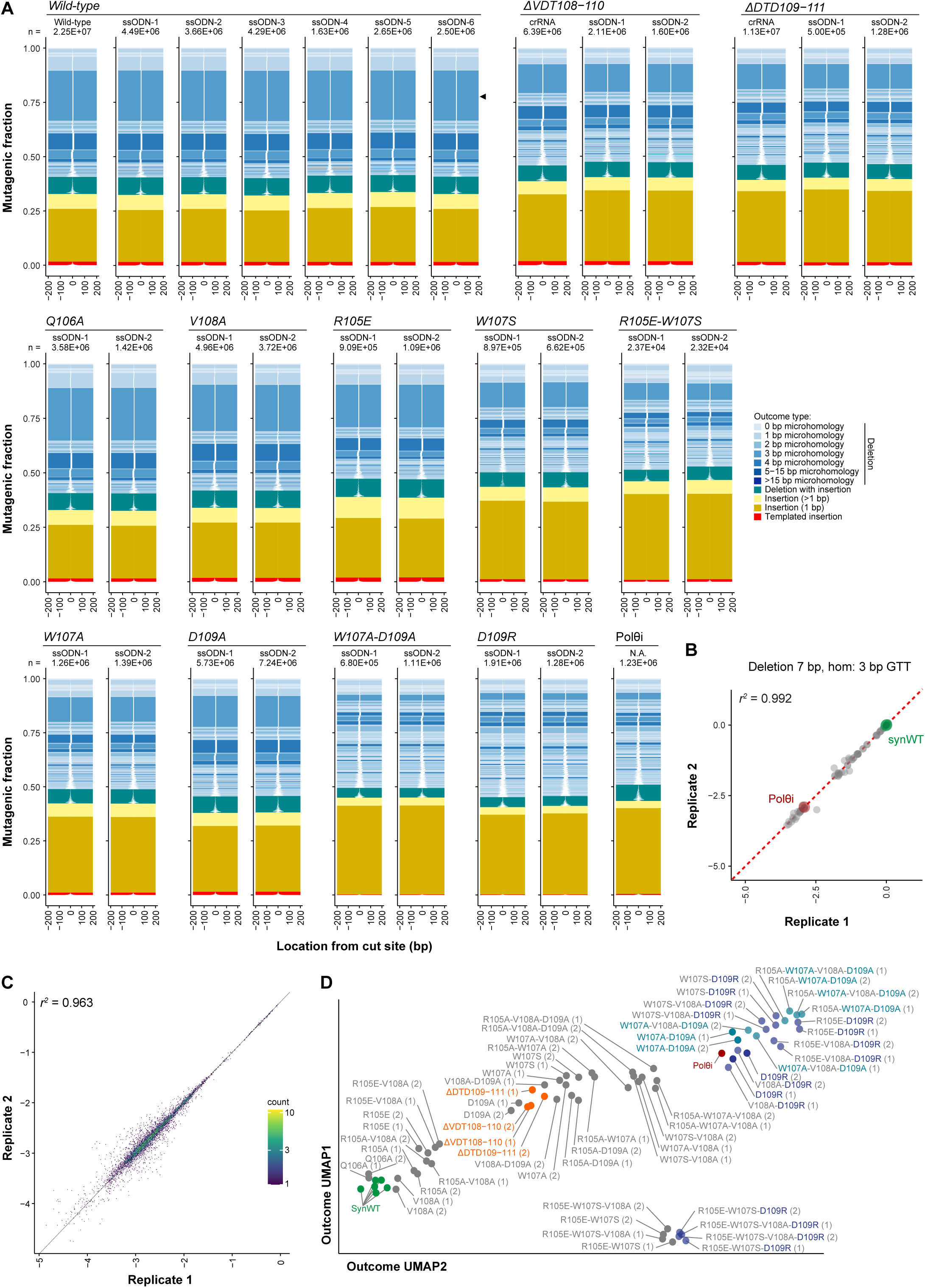
MUSIC-MAP identifies specific RPA2 substitutions that abrogate TMEJ. (**A**) Tornado plots of targeting of the *Rpa2* intron showing the complete spectrum of mutagenic repair outcomes associated with the indicated *Rpa2* alleles, as determined by MUSIC-MAP. Alleles labeled crRNA represent variants generated by Cas9 cleavage and subsequent repair, whereas ssODN indicates alleles introduced using donor oligonucleotides. Independent synonymous encodings of the same amino acid substitutions are shown separately and serve as internal controls. Individual repair products are plotted according to their frequency among mutagenic reads (y-axis) and position relative to the cleavage site (x-axis). Outcomes are colored by repair product type, with white regions indicating deleted sequences. Data from two biological replicates (independent pooled RNP transfections) were combined for visualization. The total number of mutagenic reads is indicated above each plot. A recurrent microhomology-mediated deletion is indicated (black arrowhead). (**B**) Reproducibility of MUSIC-MAP measurements across two biological replicates, showing the log2 fold-change in frequency of a representative 7 bp deletion with 3 bp microhomology (black arrowhead in (A)). Each point represents an individual *Rpa2* ssODN-produced allele variant, plotted according to its log2 fold-change in frequency. The Pearson correlation coefficient (*r^2^*) between two biological replicates is indicated. (**C**) Pearson correlation coefficient (*r^2^*) of allele-associated repair profiles between two biological replicates. Each point represents the fraction of each individual repair outcome (log-10 transformed) associated with each variant allele, shown as a density plot. (**D**) Fully annotated UMAP projection of allele-associated repair profiles shown in Fig. 1H.

**Figure S5.**
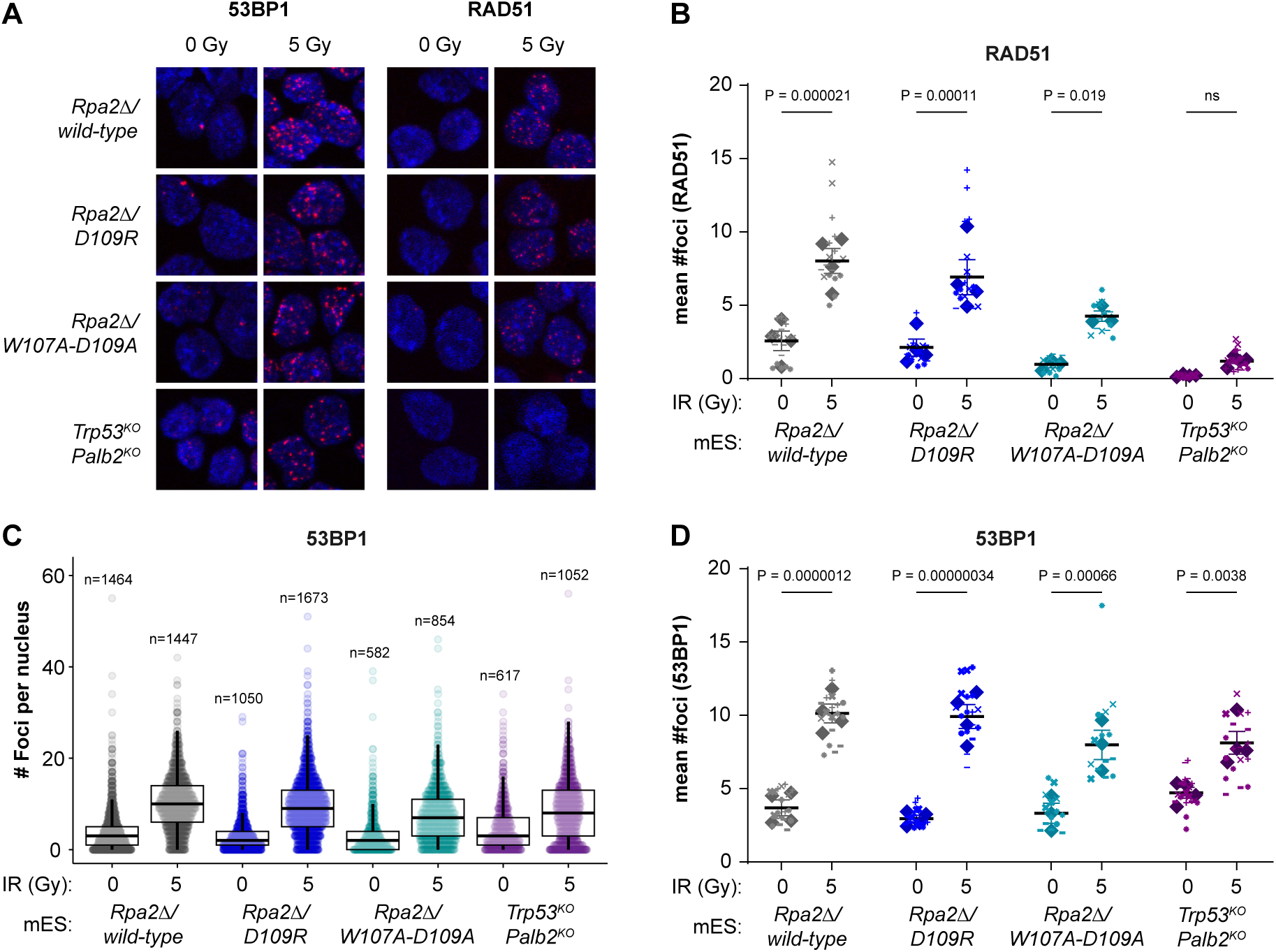
Ionizing radiation-induced RAD51 and 53BP1 focus formation in RPA2 interface mutants. (**A**) Representative confocal images of nuclei from the indicated mES cell genotypes, stained for 53BP1 or RAD51 at 3 hours after exposure to 0 or 5 Gy ionizing radiation (IR), as also presented in Fig. 3A. (**B**) Quantification of RAD51 foci, as also presented in Fig. 3A. At least four fields were imaged per genotype and biological replicate, with at least three independent biological replicates. Individual symbols (+, −, × and *) indicate the mean number of foci per field, whereas diamonds represent the mean of field averages for each biological replicate. Statistical significance was determined by CFAssay in R using ANOVA on biological replicate means, followed by Bonferroni correction for multiple comparisons. Exact P values are indicated; ns, not significant. (**C**) Box-and-whisker plots showing the distribution of 53BP1 foci per nucleus across individual nuclei pooled from all biological replicates. The total number of nuclei analyzed is indicated above each group. (**D**) Quantification of 53BP1 foci. Individual symbols indicate mean foci counts per field, and diamonds represent biological replicate means, as described in (B). Statistical analyses were performed on biological replicate means using ANOVA followed by Bonferroni correction.

**Figure S6.**
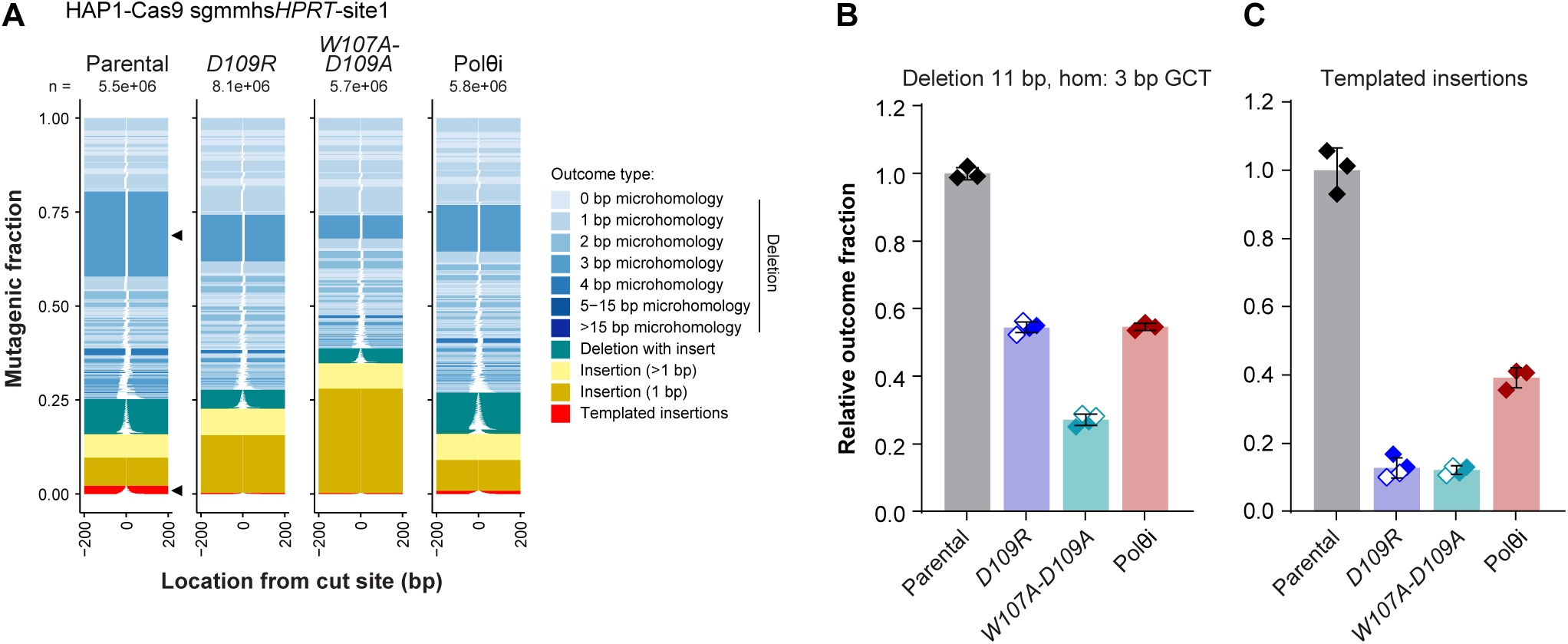
Validation of TMEJ defects in clonal HAP1 cells at an independent genomic target site. (**A**) Tornado plots showing the complete spectrum of mutagenic repair outcomes at an additional genomic target site (*HPRT*) in HAP1-Cas9 clones carrying the indicated *RPA2* mutations. Individual repair products are plotted according to their frequency among mutagenic reads (y-axis) and position relative to the cleavage site (x-axis). Outcomes are colored by repair product type, with white regions indicating deleted sequences. Biological replicates were combined for visualization, and the total number of mutagenic reads is indicated above each plot. Treatment with a Polθ inhibitor ART558 (10 µM) served as a TMEJ-deficient control. A recurrent microhomology-mediated deletion and templated insertions are indicated (black arrowheads). (**B–C**) Quantification of the relative frequencies of a Polθ-dependent 11 bp deletion with 3 bp microhomology (**B**) and templated insertions (**C**). Individual clones are represented by filled and open diamonds (biological replicates plotted separately). Bars indicate mean ± SD. Statistical significance was assessed using ordinary one-way ANOVA followed by Dunnett’s multiple-comparisons test, comparing each genotype or condition with the parental control. \*\*\*\**P* < 0.0001 (adjusted). Hom, microhomology.

**Figure S7.**
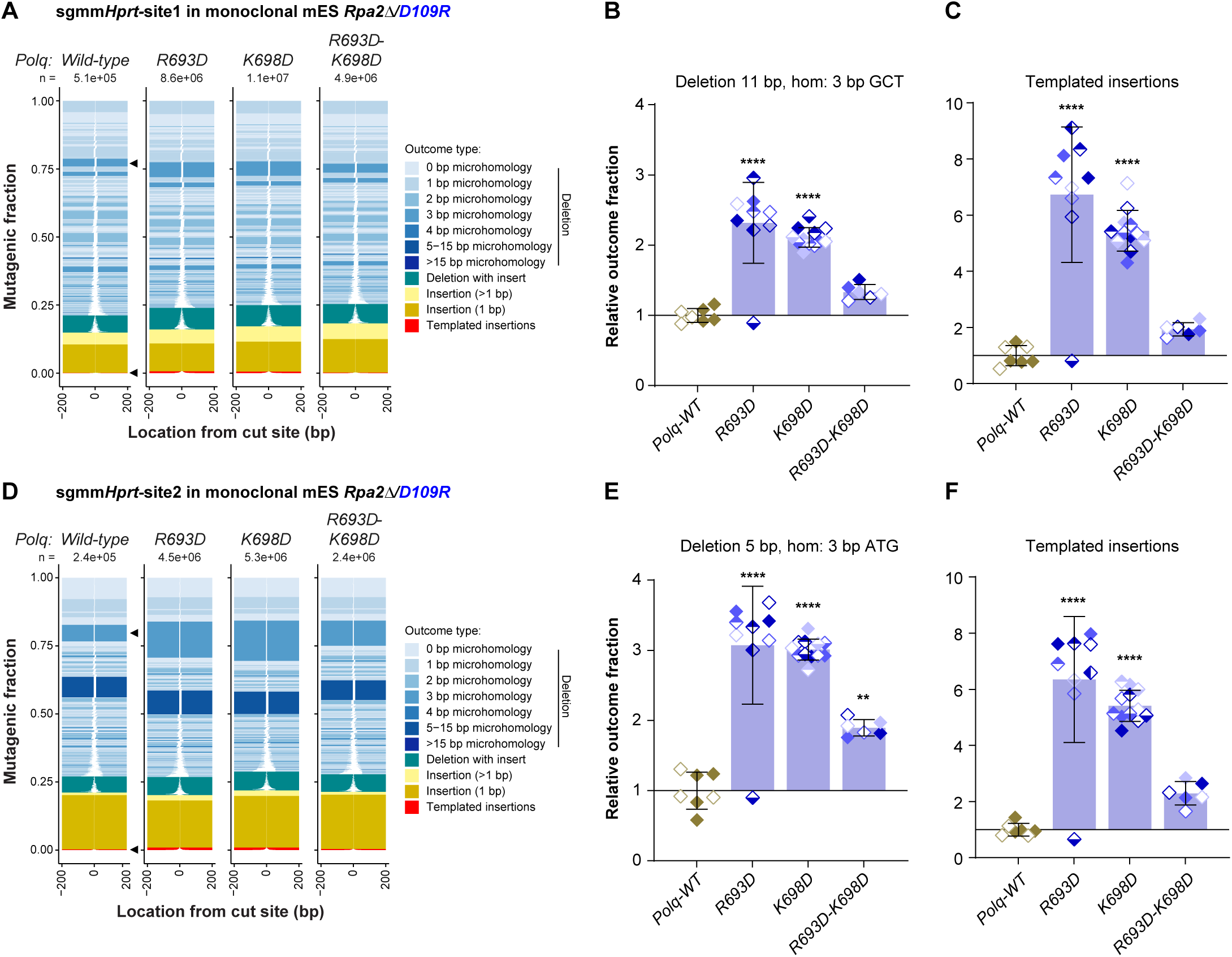
Genetic rescue of RPA2 D109R-associated TMEJ defects at two genomic target sites within the *Hprt* locus. (**A,D**) The experiment shown in Fig. 4G was repeated at two additional genomic target sites. Tornado plots showing the complete spectrum of mutagenic repair outcomes at two *Hprt* target sites in monoclonal *Rpa2* D109R mES cell lines carrying the indicated *Polq* alleles. Individual repair products are plotted according to their frequency among mutagenic reads (y-axis) and position relative to the cleavage site (x-axis). Outcomes are colored by repair product type, with white regions indicating deleted sequences. Repair profiles from independent clones of the same genotype were combined for visualization, and the total number of mutagenic reads is indicated above each plot. Recurrent microhomology-mediated deletions and templated insertions are indicated (black arrowheads). (**B,C,E,F**) Quantification of the relative frequencies of a recurrent 11 bp deletion with 3 bp microhomology (**B**), templated insertions (**C**), a recurrent 5 bp deletion with 3 bp microhomology (**E**), and templated insertions (**F**), as indicated by black arrowheads in (A,D). Individual clones are represented by different symbols (biological replicates shaded separately). Outcome fractions were normalized to those of the parental *Rpa2 D109R* clone carrying wild-type *Polq*. Bars indicate mean ± SD. Statistical significance was assessed using ordinary one-way ANOVA followed by Dunnett’s multiple-comparisons test, comparing each genotype with the parental control. \*\**P* < 0.01, \*\*\*\**P* < 0.0001 (adjusted). Hom, microhomology.

**Figure S8.**
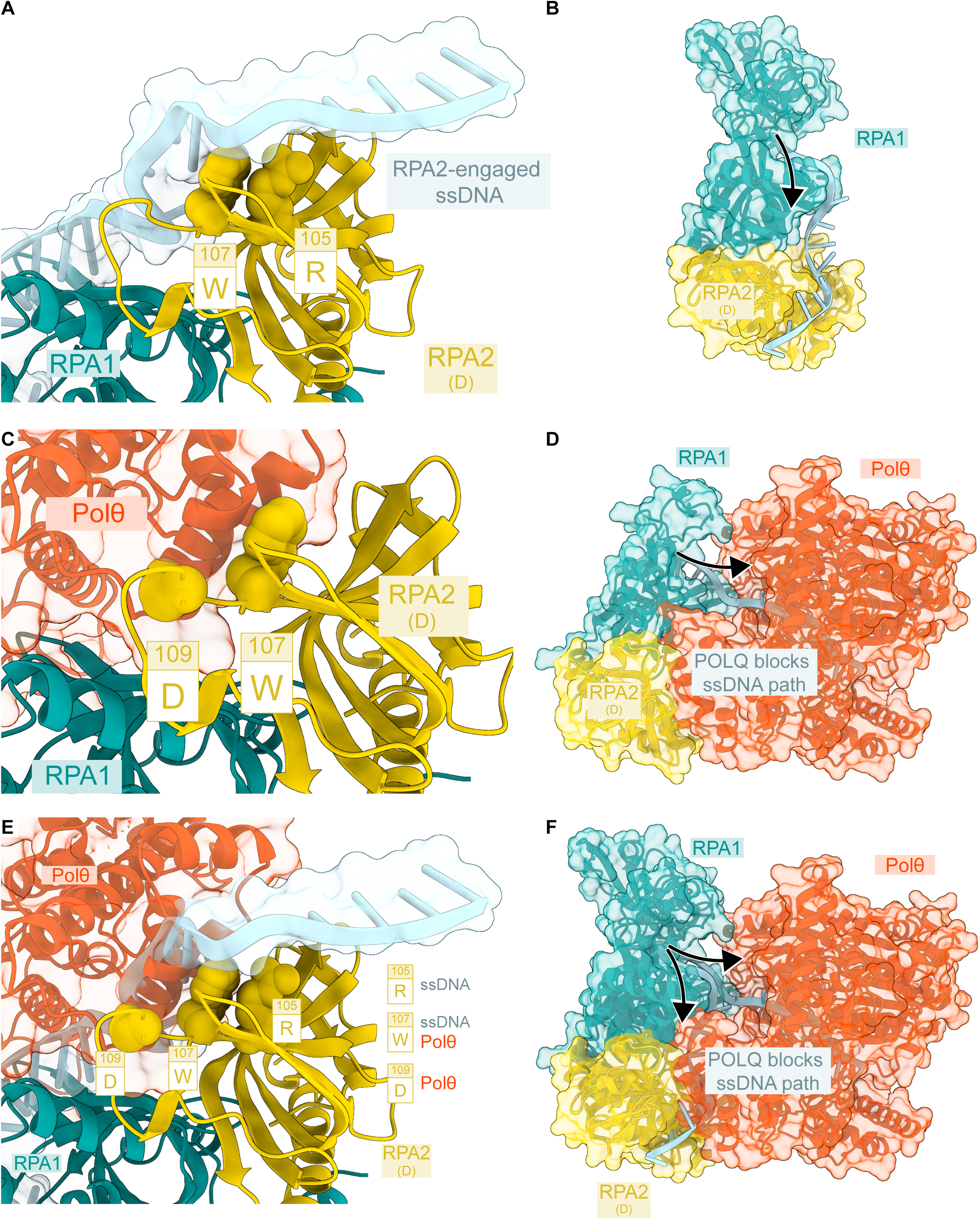
AlphaFold3-based structural modeling suggests how Polθ engages RPA-coated DNA. (**A–F**) Structural comparison of AlphaFold3 models of RPA (mouse) bound to ssDNA and RPA (mouse) associated with Polθ (mouse). (**A,B**) Model of the RPA1-RPA2 complex bound to ssDNA, highlighting the positioning of ssDNA along the RPA2 DNA-binding domain D (DBD-D) and the proximity of RPA2 residues R105 and W107 to the ssDNA-binding surface. (**C,D**) Model of the RPA1-RPA2 complex associated with the Polθ N-terminal helicase domain, showing the predicted interaction involving RPA2 residues W107 and D109. In this configuration, Polθ occupies a position that would interfere with the trajectory of ssDNA through the RPA complex. (**E,F**) Structural overlay of the two models, illustrating the spatial relationship between the predicted Polθ-binding interface and the ssDNA-binding path. The superposition reveals a potential steric incompatibility between Polθ binding and the modeled trajectory of ssDNA along the RPA complex.

## Notes

### Competing Interest Statement

The authors have declared no competing interest.

